# MAGPIE: an interactive tool for visualizing and analyzing protein-ligand interactions

**DOI:** 10.1101/2023.09.04.556273

**Authors:** Daniel C. Pineda Rodriguez, Kyle C. Weber, Belen Sundberg, Anum Glasgow

## Abstract

Quantitative tools to compile and analyze biomolecular interactions among chemically diverse binding partners would improve therapeutics design and aid in the study of molecular evolution. Here we present MAGPIE (Mapping Areas of Genetic Parsimony In Epitopes), a publicly available software package for simultaneously visualizing and analyzing thousands of interactions between a single protein or small molecule ligand (the “target”) and all of its protein binding partners (“binders”). MAGPIE generates an interactive 3D visualization from a set of protein complex structures that share the target ligand, as well as sequence logo-style amino acid frequency graphs that show all the amino acids from the set of protein binders that interact with user-defined target ligand positions or chemical groups. MAGPIE highlights all the salt bridge and hydrogen bond interactions made by the target in the visualization and as separate amino acid frequency graphs. Finally, MAGPIE collates the most common target-binder interactions as a list of “hotspots,” which can be used to analyze trends or guide the *de novo* design of protein binders. As an example of the utility of the program, we used MAGPIE to probe how two ligands bind orthologs of a well-conserved glycolytic enzyme for a detailed understanding of evolutionarily conserved interactions involved in its activation and inhibition. MAGPIE is implemented in Python 3 and freely available at https://github.com/glasgowlab/MAGPIE, along with sample datasets, usage examples, and helper scripts to prepare input structures.

## 1. Introduction

It is challenging to determine patterns in a large set of biochemically diverse protein-ligand interactions. For example, analyzing how a library of computationally designed proteins interact with a binding partner usually requires ranking the design models by several quality metrics, and then inspecting top-ranked models in detail individually. Similarly, although assembling a multiple sequence alignment (MSA) of hundreds of evolutionarily related proteins is expedited by bioinformatics tools, it remains difficult to visualize how sequence variability at one or more positions in a ligand binding pocket manifests in three dimensions (3D). The problem is exacerbated when important protein regions are non-contiguous in sequence space and when the protein can interact with multiple binding partners. Thus, there is a need for a versatile protein complex analysis tool that represents the sequence diversity and biochemistry inherent in molecular interactions on the fly.

A method to identify sequence conservation and variability among proteins that bind to common ligands in 3D would help us understand the biochemical requirements for key protein interactions and functions and engineer new proteins that respect these requirements. However, existing methods only partially address this need. For example, using sequence logos to study protein-protein interactions requires that all the protein binder sequences are similar enough to align in an MSA.^1^ MSA generation algorithms present limitations for investigating protein interactions in 3D as they are designed for aligning sequences of evolutionarily related proteins with some sequence identity. Further, MSAs cannot be used to align small molecule ligands or map their interactions with proteins.^2–7^ Small molecule alignment methods are primarily implemented as computer-aided drug discovery (CADD) software for determining native binding poses or improving docking for pharmacological applications, but these do not typically include methods for analyzing biochemical interactions and can be difficult to use in the context of protein complexes.^8–13^ And while protein structural search methods such as TM-align^14^, Dali^15^, and FoldSeek^16^ have enabled the identification of proteins with local structurally homologous regions, and programs like SSDraw can align sequences to show how secondary structures are conserved in a set of homologous proteins^17^, these methods are not designed to highlight or organize conserved binding interactions across diverse proteins: for example, small molecule ligands that bind multiple proteins via different binding pocket geometries or antigens that bind a library of engineered antibodies at non-overlapping epitopes. In such cases, translating trends at the 2D sequence level to the 3D space towards identifying molecular patterns that drive or tune biochemical interactions presents a challenge for which no generalizable software is available, even when high-confidence interactions can be predicted at a large scale.^18–23^

To meet this growing need in protein science, and complementing recent methods for visualizing protein-peptide interactions^24^ and protein structural features^25^, we introduce MAGPIE: Mapping Areas of Genetic Parsimony In Epitopes. MAGPIE is a protein complex visualization and analysis software to facilitate the simultaneous comparison of many protein-partner interactions in 3D by structure and sequence. MAGPIE identifies amino acids (AAs) in a set of proteins that make molecular interactions with a user-defined region on a single target ligand, which can be a protein or a small molecule. MAGPIE also generates AA frequency graphs on the fly that show how often different AAs in the set of proteins interact with specific target ligand AAs or heavy atoms (HAs). Furthermore, MAGPIE generates an interactive 3D representation of the target ligand and visually illustrates the positions of all alpha carbons (Cα) that are near it, coloring these by AA biochemical characteristics, highlighting patches of conserved or similar interactions, and collating trends as a list of “hotspots” that can be used to inform the computational design of new protein binders. We anticipate that MAGPIE will be useful in applications ranging from protein design, where it can identify biases in computational methods for engineering protein-ligand interactions, to molecular biology, where it could illuminate functional adaptation in proteins by revealing how their features have emerged and changed over evolutionary history.

## 2. Results

### 2.1. Software overview

MAGPIE operates in several steps in a Jupyter notebook format with optional helper scripts in Python for input dataset preparation (Figure 1). As inputs, the user provides a set of protein complex structures in Protein DataBank (PDB) format that are structurally aligned on the target ligand (Figure 1A). The target ligand must have the same chain index in all input structures, and the protein binding partners must also share a chain index that is different from the target ligands. The helper scripts aid in preparing datasets for MAGPIE by automatically renumbering atoms, renaming chains, protonating protein residues, and aligning the structural models on the target ligands using either a reference structure from the dataset or an MSA (Figure 1A, B). For small molecule targets, the alignment helper script generates separate sets of aligned structures for different ligand conformational isomers (“conformers”) with a user-specified global root-mean-square deviation (RMSD) threshold (“conformer pools”) (Figure 1B). MAGPIE records the positions of all atoms within the target chain in all the structures. Subsequently, it also stores the positions of all atoms from the binder chains in the dataset (Figure 1C, i).

**Figure 1.**
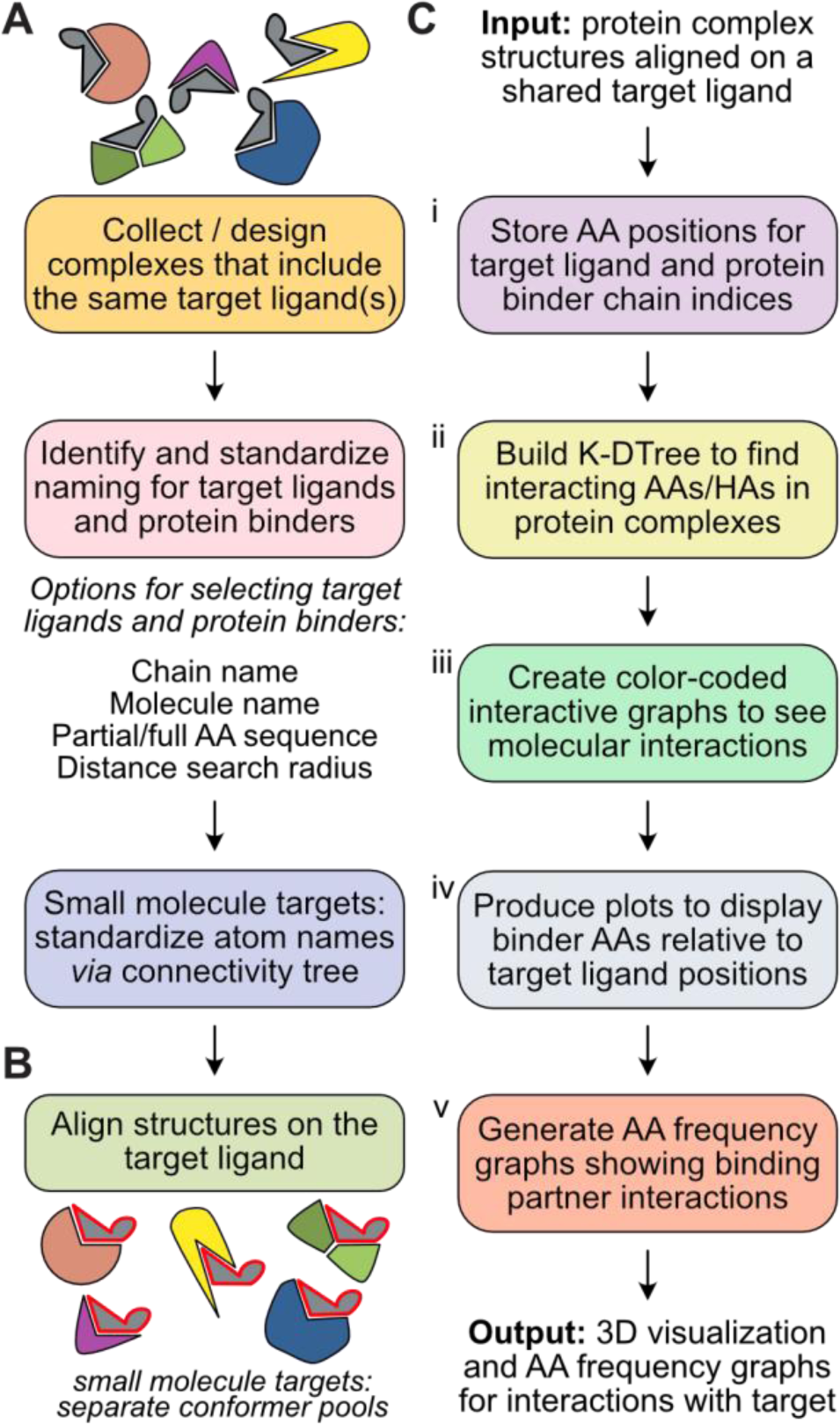
MAGPIE framework. **(A)** The user compiles a set of protein complex structural models that share a target ligand, which can be a small molecule or a protein. The helper script MAPIE_input_prep.py cleans and processes the models. **(B)** The user can align the models on the target ligand using the helper scripts. The small molecule alignment script sorts the models into different conformer pools. **(C)** MAGPIE produces an interactive graph of protein binder interactions with amino acids (AAs) and heavy atoms (HAs) on the target ligand, which can be toggled to show hydrogen bond and salt bridge interactions, as well as distance-defined hotspots colored by AA biochemical characteristics. MAGPIE also generates AA frequency graphs showing binder interactions with user-defined target AAs or HAs.

Second, MAGPIE employs a K-DTree to efficiently query which AAs from the binding partners fall within a user-defined distance in Ångstroms (Å) of any AAs/heavy atoms (HAs) from any atoms in the target ligand (Figure 1C, ii).^26^ This step identifies potential binding partner AA interactions with the target ligand. MAGPIE calculates distances and torsion angles for all binder AAs in this group with respect to the nearest target AAs/HAs to identify hydrogen bond and salt bridge interactions and to determine trends across the set of protein binders.

Third, MAGPIE employs the Plotly library to create a color-coded interactive 3D visualization, displaying the 3D locations of all binding partner residue Cα and the target ligand structure main chain or full molecular structure (Figure 1C, iii).^27^ This allows users to explore the spatial arrangement of the protein-ligand interactions by zooming in on specific regions of the complex, panning and rotating the structure, and hovering over target and binding partner AAs/HAs to see their identities and visually spot trends in the interactions. The target structure is represented in black (for protein targets) or CPK coloring (for small molecules), while the binder AAs can be colored using the RasMol-based “Shapely” or “Amino” colors, which groups them by their biochemical characteristics, such as AA charge, hydrophobicity, and size.^28^

MAGPIE then generates plots to display the frequency of AAs among the binder residues within a user-specified distance from target positions, which can be selected and changed by the user *ad hoc* while exploring the 3D visual representation (Figure 1C, iv). MAGPIE includes options to customize a subset of the target ligand AAs (for protein ligands) or HAs (for small molecule ligands) for detailed analysis in specific regions. Once the positions are selected, MAGPIE identifies binding partner residue Cα atoms using the K-DTree from Step 2, calculates the frequency of each AA in the set of binders within the user-specified distance, and generates an AA frequency graph using the LogoMaker Python package (Figure 1C, v).^29^ The AA frequency graph shows the HA or AA index position in the target ligand *vs.* the frequency of AAs among binding partner residues from the input PDB dataset found within the specified distance. Each one-letter AA code is represented by a height that corresponds to its frequency in the set of binding partners. The AA frequency graphs help to quantify the distribution and conservation of AAs at specific positions involved in the binding interactions with the target. The number of individual interactions counted for each column of the graph is reported in the *x* axis label. Further, the AA frequency graphs can be customized to show hydrogen bond or salt bridge partners for a quantitative readout of specific types of interactions at different target positions in the dataset. In the 3D visualization, the colors of the protein binder Cα can be toggled to show hydrogen bond or salt bridge partners in red, with all other AAs in white.

To collate and report trends in interactions with the shared target ligand, MAGPIE additionally produces a data table of ligand AAs or HAs that participate in hydrogen bonds and salt bridges that are enriched in the set of protein binders. MAGPIE uses the DBSCAN algorithm implemented in Python Scikit-learn to generate a list of “hotspots” to identify the AAs that are involved in the most prevalent interactions with the target ligand and group them by their biochemical characteristics.^30,31^ The hotspots can be used to guide efforts in reengineering important protein-ligand interactions or build protein binding partners *de novo* using generative design methods.^32,33^ The parameters for identifying hotspots, such as how many AA are required to form a hotspot and how far away they may be from each other in Cartesian space, are also user-adjustable. The detailed MAGPIE methods and algorithms are available in section 2 of the supplemental material.

We present three case studies to demonstrate how MAGPIE can be used. First, we explore trends in how dozens of antibodies bind to the same target antigen (Figures 2 and 3). Second, we study how a common small molecule metabolite interacts with sequence-and structure-diverse protein binding partners in a large set of natural complexes (Figure 4). Finally, we build and analyze structural models of distantly related bacterial orthologs of a central glycolytic enzyme bound to allosteric ligands (Figures 5 and 6).

**Figure 2.**
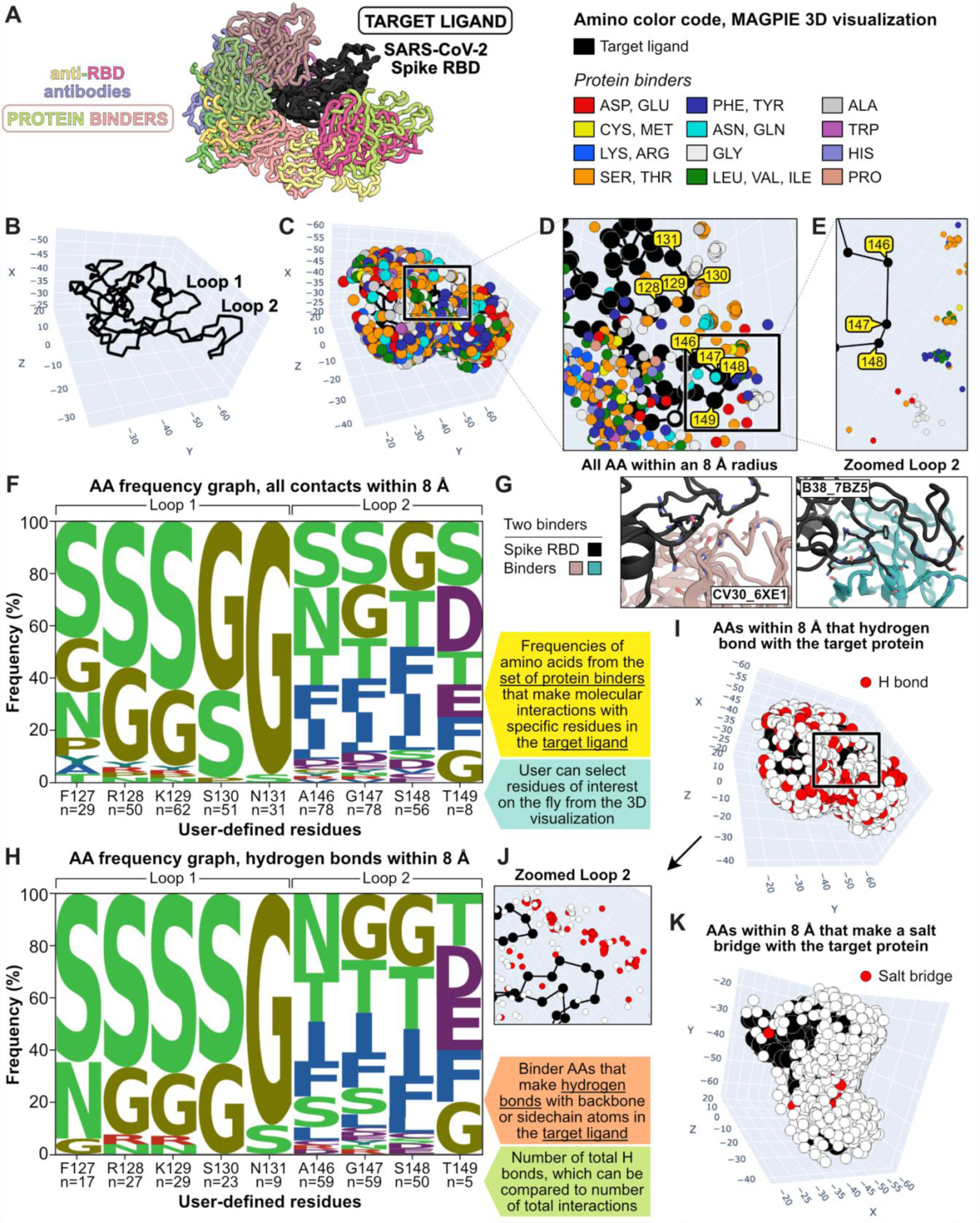
Protein-protein interactions: SARS-CoV-2 spike receptor binding domain binds a variety of antibody fragments. **(A)** Structural representation of the SARS-CoV-2 spike receptor binding domain (RBD, black) bound to 15 anti-RBD binders (colors), from a set of 63 binders. **(B)** MAGPIE representation of the protein target main chain with two loops labeled as an epitope of interest. **(C-E)** MAGPIE representations of the RBD-binder complexes showing all AA from the binders within 8 Å of the RBD epitope. The RBD Cα are shown as black circles and the anti-RBD binder Cα are shown as colored circles using the Amino color code as shown. The binder AAs are distinguishable in 3D space as the user zooms in. **(F)** AA frequency graph quantifying all neighboring AA for several RBD positions in both loops, as shown in the 3D visualization. The number of interactions and the RBD positions are listed in the *x*-axis. **(G)** Structural models of two antibodies that bind the epitope of interest. Loop 1 interacts with serines and glycines in the antibodies in both examples, while larger hydrophobic residues are enriched in Loop 2. **(H)** AA frequency graph showing hydrogen bond partners for every position specified in the epitope. **(I)** 3D MAGPIE visualization of H bond partners within 8 Å of the RBD epitope. **(J)** Zoomed in Loop 2 H bond partners. **(K)** The 3D visualization can also highlight AAs in the binders that make salt bridges with the target ligand.

**Figure 3.**
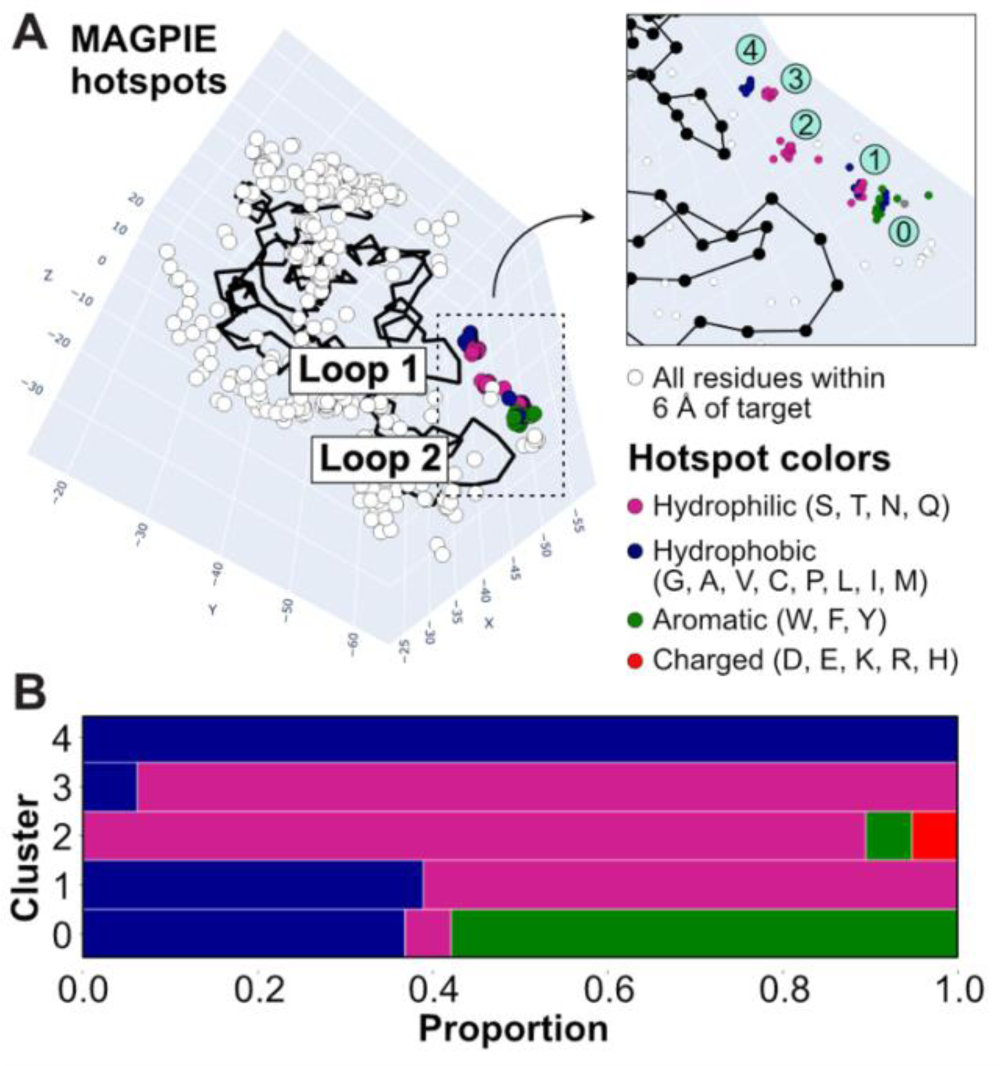
Binding hotspots. **(A)** Default settings in MAGPIE’s hotspot finding feature revealed that binder AAs neighboring two loops in the RBD organize into **(B)** five hotspot clusters.

**Figure 4.**
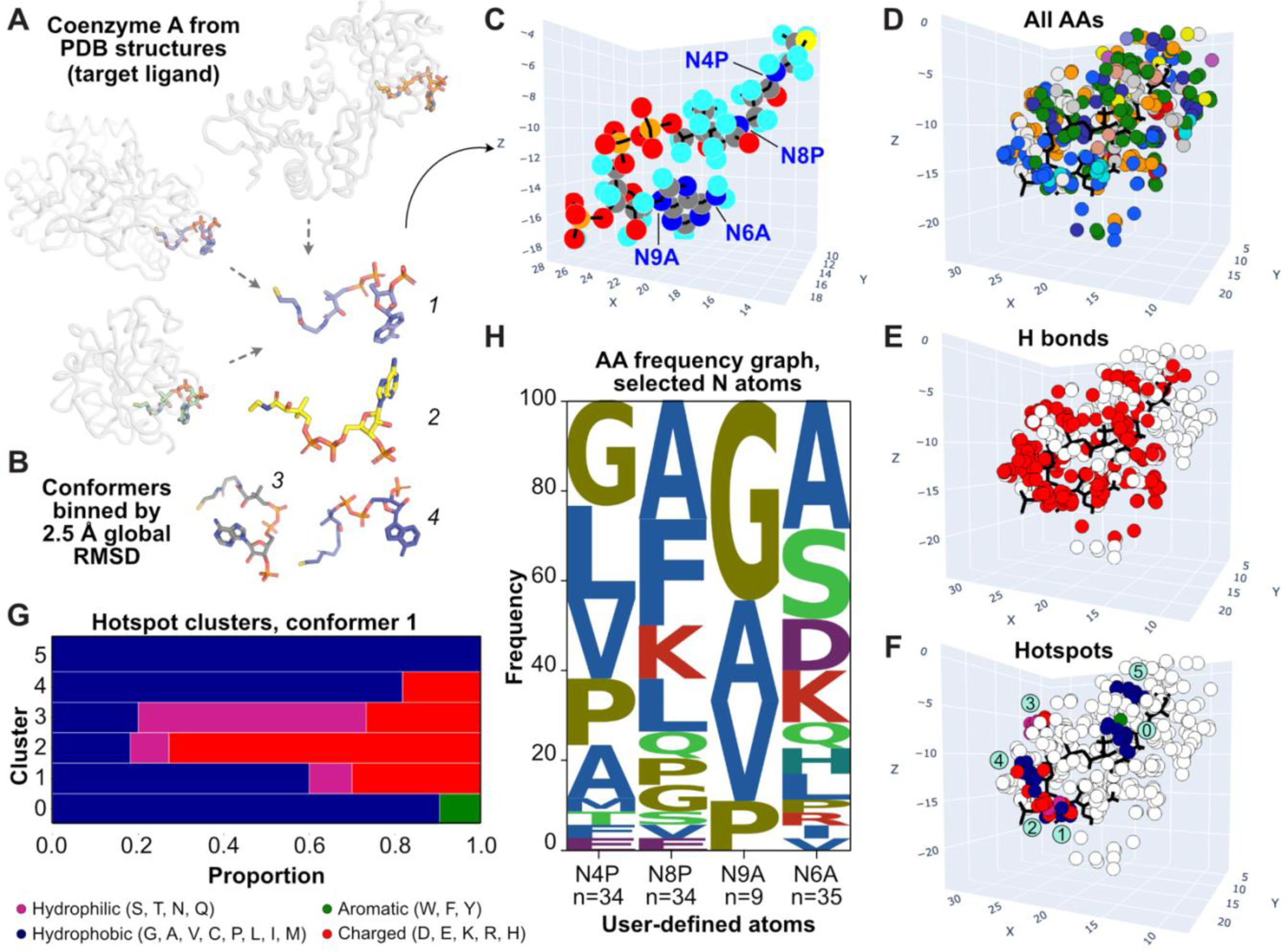
Protein-small molecule interactions: coenzyme A binds diverse proteins. **(A)** A subset of 199 complexes were randomly chosen from 608 CoA-protein complexes in the PDB and sorted into 31 conformer pools. **(B)** Examples of conformer structures shown as sticks. **(C)** CoA conformer 1 shown as a ball-and-stick model in the MAGPIE 3D visualization. Conformer Pool 1 included 22 structures. **(D-F)** Binder AAs within 5 Å of CoA in Conformer Pool 1: (D) all interaction partners colored by the Amino code; (E) H bond partners, red; (F, G) hotspots. (H) AA frequency graph showing nearest neighbors for CoA Conformer 1 atoms labeled in (C).

**Figure 5.**
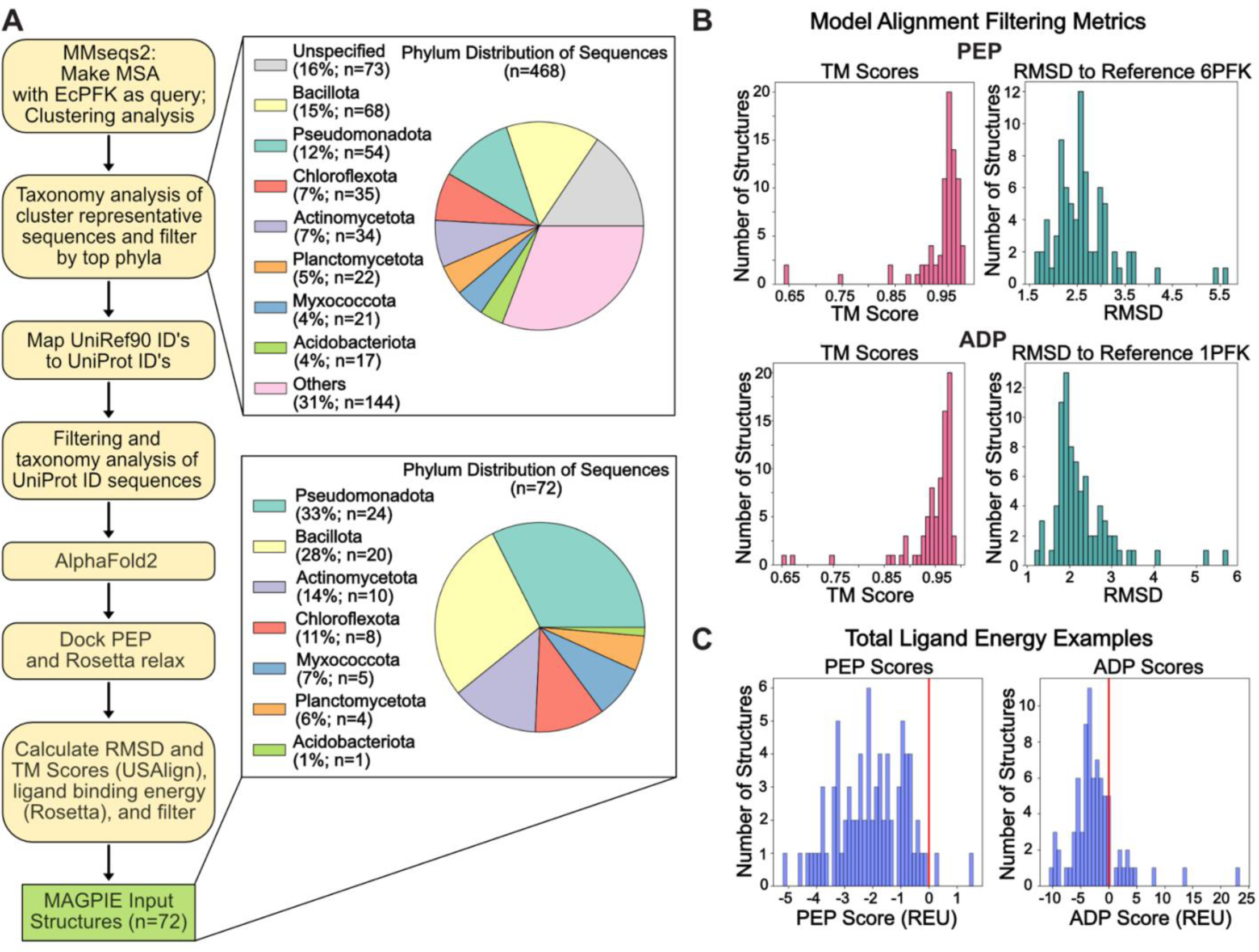
MAGPIE input structure preparation pipeline for case study 3. **(A)** MAGPIE input structures were prepared and filtered using a structural bioinformatics and modeling pipeline. Phylum distribution at various filtering steps is shown in the pie charts. **(B)** Distribution of RMSD and TM-score calculated by US-align for PEP and ADP. **(C)** Rosetta-calculated ligand energies for PEP- and ADP-bound bacterial PFK-1 models used for a final filtering step of MAGPIE input structures. Structures with a positive ligand energy were excluded.

**Figure 6.**
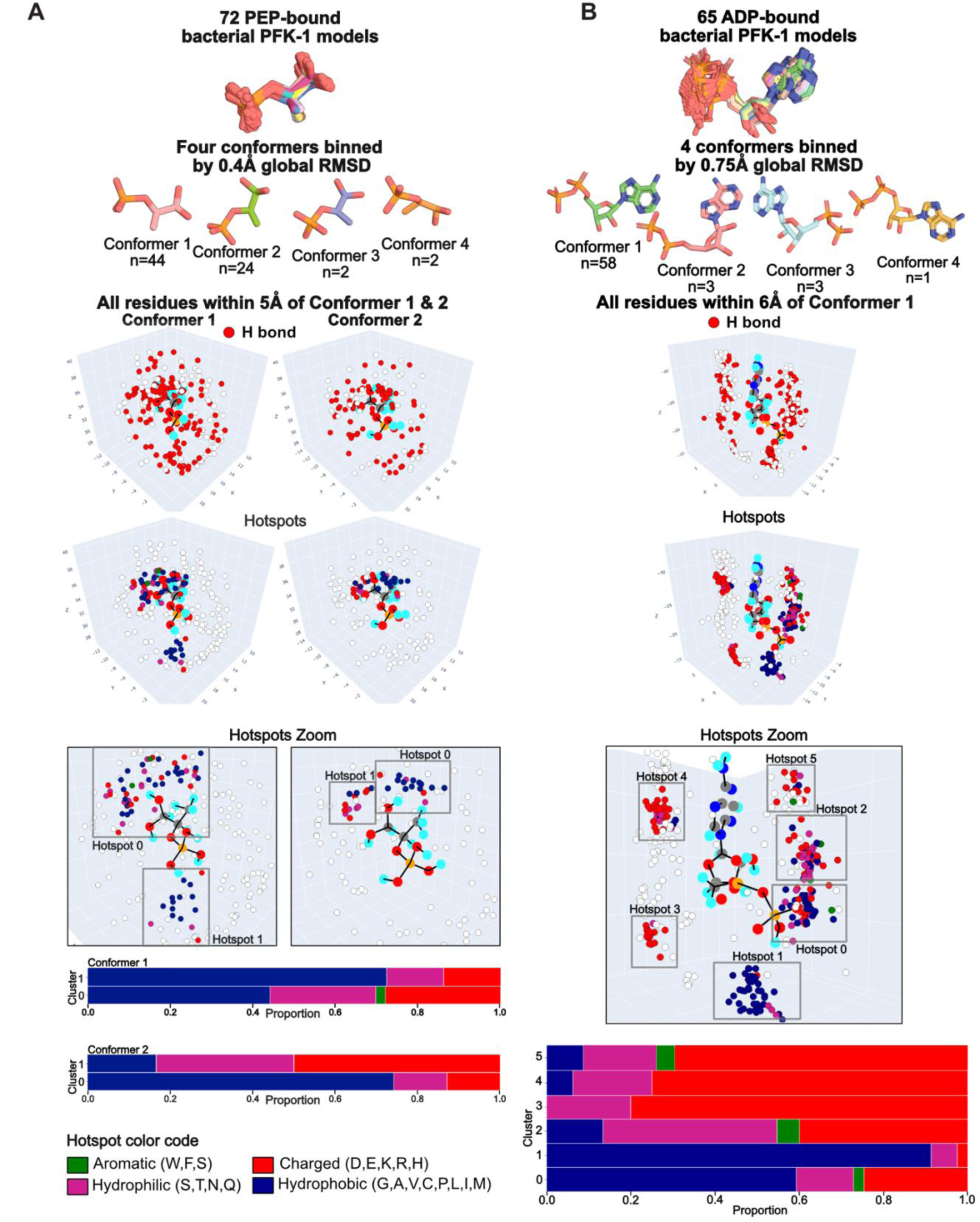
MAGPIE analysis of ligand-protein binding interactions of the allosteric effectors of PFK-1. **(A)** Feeding 72 PEP-bound bacterial PFK-1 models into MAGPIE resulted in four PEP conformer pools with similar hydrogen bonding partners and hotspots enriched in AAs with different biochemical characteristics. **(B)** 65 ADP-bound bacterial PFK-1 structures were separated into four conformer pools. Hydrogen bonding partners and hotspots are shown for the dominant conformer.

### 2.2. Case study 1: antibody-antigen complexes in which the antigen is the target ligand

Figure 2 illustrates how MAGPIE can be used to study protein-protein interactions. In this case study, we explored how a set of 63 nonredundant antibodies (specifically their fragment antigen-binding regions, or Fabs) and nanobodies bind to the SARS-CoV-2 spike protein receptor binding domain (RBD). A subset of these binders are shown aligned on the RBD in Figure 2A.^34^ MAGPIE aids in visualizing the complete collection of antibody- and nanobody-binding epitopes within 8 Å of the RBD, with the binder paratope AAs colored according to the Amino color code (Figure 2B, C). MAGPIE shows which RBD epitopes are most involved in these interactions, as well as the prevalence of different binder AAs, as the user zooms, pans, and toggles among coloring schemes that highlight all interactions, hydrogen bonds, and salt bridges. We focused on Fab and nanobody interactions with a common RBD epitope comprising two loops (Figure 2B). Of the 63 binders, 27 interact with at least one of the two loops in this region via their complementarity-determining regions (CDRs) (Figure S1).

MAGPIE compiles all the models in one place and allows the user to see the local environment around every AA in the two loops at different levels of detail and from any angle (Figure 2C-E). We generated an AA frequency graph to show which antibody and nanobody AAs are enriched in the local vicinity of every position in this RBD epitope in the full dataset (Figure 2F). MAGPIE revealed that the two loops in the epitope participate in markedly different interactions: while the first loop (residues 127-131) interacts almost exclusively with serines and glycines in this set of binders, the AAs that interact with the second loop (residues 146-149) are more diverse and include polar, hydrophobic, and charged side chains. This information allows the user to focus on specific target residues when inspecting individual complexes in the dataset to understand the molecular details of each enriched interaction more deeply (Figure 2G).

MAGPIE further revealed that RBD AAs in this epitope frequently hydrogen bond with the binders in the dataset (Figure 2H-J). The hydrogen bond AA frequency graph shows that in the first loop, serines account for 60% of the hydrogen bond partners, while glycine backbone interactions make up most of the remaining hydrogen bonds. The enrichment of these small AAs in the loop suggests close proximity of the CDR backbone to the RBD in this region (Figure 2H). The hydrogen bonding partners in the second RBD loop in the AA frequency graph are more diverse and include larger AAs that are both polar and hydrophobic, suggesting that the small hydrophobic residues A146 and G147 at the beginning of this loop form backbone hydrogen bonds and hydrophobic interactions with the binders. Closer inspection of structures in the dataset confirms this hypothesis. For example, in Figure 2G, the small hydrophobic AAs at positions 146 and 147 in the second loop make backbone hydrogen bonds as well as close-packed hydrophobic interactions with one antibody CDR, whereas the other loop is involved in backbone and side chain hydrogen bonds with serines spanning multiple CDRs.

We observed that these RBD AAs make additional nonpolar interactions with hydrophobic and charged residues in the binders, which is suggested by the visualization (Figure 2D) and confirmed by comparing the AA frequency graphs for all contacts vs. hydrogen bond partners only (Figures 2F, H). For example, about 15% of interactions with F127 in the first loop are with hydrophobic AAs proline, tyrosine, and alanine, but none of these AAs contribute to any hydrogen bonds observed with F127 in the dataset. However, F127 pi stacks with these AAs in individual structures (Figure 2G). MAGPIE can also identify salt bridges in protein-protein complexes (Figure 2K), though we found that this two-loop epitope does not participate in any. The user has the option to output a .csv file that contains all the detailed information about every hydrogen bond and charged interaction observed in the pool.

Finally, towards identifying optimal epitopes for therapeutic targets, analyzing docked models, or choosing positions for *de novo*-designed binding interactions, MAGPIE can identify hotspots: groups of AAs in the pooled binding partners that are close in 3D space and have similar biochemical characteristics (Fig. 3). The distance between AAs that constitute a hotspot and the number of AAs required for a hotspot are adjustable by the user in the hotspot selection feature. Using the default parameters of a 2 Å distance threshold and 15 AA minimum to identify the hotspots, MAGPIE identified five hotspots in this region with varying biochemical characteristics. The biochemical makeup of the AAs in each hotspot are summarized in a graph output (Fig. 3), with a simplified color scheme similar to the Amino scheme in the full 3D MAGPIE visualization. MAGPIE also outputs a .csv file that contains the hotspot residues and their exact AA composition.

In summary, the AA frequency graphs generated by MAGPIE showcase the diversity of interactions with Fabs and nanobodies from the SARS-CoV-2 spike RBD binder dataset for any user-defined AAs in the RBD belonging to any region, which can be altered *ad hoc*. Further, MAGPIE identification of CDR biochemical features that we observed to facilitate binding to the RBD in this dataset could enable the rapid design of antibody-alternative protein therapeutics that bind the antigen with a large, highly structured binding interface^35–38^, which may prove to be more robust to viral mutations than a flexible antibody CDR.

### 2.3. Case study 2: protein complexes with a shared small molecule target ligand

MAGPIE can also be applied to study protein-small molecule complexes. For example, identifying trends in how a specific small molecule binds structurally diverse proteins could provide guidance for designing a binding partner *de novo* for a chemically similar ligand.^39,40^ The common metabolite coenzyme A (COA) binds hundreds of different proteins from a variety of organisms, and experimentally solved structures are available for many of these complexes in the PDB. Additionally, COA presents a challenging target for designing new protein binders given its inherent flexibility, with four main rotatable bonds that aid in its function as a carrier of acyl groups in various metabolic reactions. It is unknown to what extent topologically diverse binding partners converge on specific shared interactions with individual functional groups in common metabolites like COA.

In our case study, MAGPIE highlighted similarities and differences among COA interactions in 199 binding pockets and interfaces from bacterial enzymes, including ligases, acetyltransferases, synthases, and epimerases (Figure 4A). We compiled the COA dataset by searching the PDB for COA-containing structural models, which yielded more than 600 hits (details in section 8 of the supplementary material). We randomly selected 199 of these and used our cleaning and alignment method (Figure 1A, B) to standardize and separate the COA-bound structures into 31 conformer pools based on a 2.5 Å RMSD threshold (Figure S2).

Representative COA structures from four conformer pools are shown in Figure 4B. MAGPIE can be used to visualize the local chemical environments for each atom in each COA conformer (Figure 4C, D, Figure S3A, B). Focusing on the first conformer pool, which includes 22 structures, we observed that several of the COA nitrogen atoms are neighbored by a combination of hydrophobic AAs and a few charged AAs (Figure 4D, H), many of which make hydrogen bonds with COA (Figure 4E). Of the nitrogen HAs in the conformer pool, only the NH_2_ group of the adenine ring (N6A, Figure 4C) interacts appreciably with polar and charged AAs.

Using a 2 Å threshold for distance among hotspot AAs and a minimum number of 20 AAs to form a hotspot, MAGPIE identifies six hotspots (Figure 4F), four of which cluster near the other nitrogen atoms in Figure 4C. The hotspots primarily contain hydrophobic and charged residues (Figure 4G). However, Hotspot 3, which is near the diphosphate group of COA and distal to any of the nitrogen atoms, includes several hydrophilic AAs.

MAGPIE can be used to investigate any COA conformer pool with more than a few members to discover how the local environment of specific functional groups changes in response to changes in the global conformation of the metabolite when it is complexed with different proteins. To probe this, we compared the COA nitrogen atoms from Conformer 1 as shown in Figure 4 to the same nitrogen atoms in the 21 structural models comprising Conformer Pool 2 (Figure S3A). Using the same MAGPIE parameters, we observed that the COA-binding proteins in this pool favor interactions that are more localized to one side of the COA molecule in the concave area between the 3’ phosphoadenosine and the cysteamine group (Figure S3B), and a smaller proportion of these interactions are with hydrophobic AAs, as compared with Conformer Pool 1 (Figure S3F). Despite including a comparable number of models as the first pool, MAGPIE finds only two hotspots in the second conformer pool, which are dominated by hydrophilic and charged AAs (Figure S3D, E). One of the hotspots neighbors the β-alanine and cysteamine nitrogens in COA, as observed in the first conformer pool (N4P, N8P, respectively). Comparing the local interaction partners for N4P and N8P between the two conformer pools, we find that they are quite different, with several charged and polar AAs enriched in the local environment in Conformer Pool 2 (Figure S3F). For example, whereas N4P in Conformer 1 is frequently supported by a nearby small or medium-sized hydrophobic residue (Figure S4), in Conformer 2 this atom tends to form hydrogen bonds with serines, asparagines, or glutamic acids (Figure S3G). The other nitrogen atoms we considered have more similar chemical environments in the two conformer pools: N9A is neighbored by valines and glycines, and N6A has a varied environment. This analysis illustrates how the flexible COA molecule might be constrained to a specific geometry in a binding site for a forward-design application.

### 2.4. Case study 3: trends in inhibitor binding for an evolutionarily conserved kinase

As a final case study involving protein-small molecule complexes, we explored how a well-conserved glycolytic enzyme, phosphofructokinase-1 (PFK-1), evolved across bacterial phyla while maintaining binding to an inhibitor and an activator in an allosteric pocket. PFK-1 performs the ATP-catalyzed conversion of fructose-6-phosphate (F6P) to fructose-1,6-bisphosphate (FBP). All known PFK-1 are allosterically inhibited by phosphoenolpyruvate (PEP) and allosterically activated by adenosine diphosphate (ADP), which bind the same allosteric site in the enzyme. We sought to determine how distantly related bacterial PFK-1 adapted to accommodate binding both PEP and ADP while maintaining their opposite regulatory effects, despite low apparent sequence identity in the allosteric pocket due to evolutionary drift.

Because few bacterial PFK-1 structures have been experimentally solved^41–45^, we instead compiled a diverse set of sequences from the Uniprot database as inputs for structural prediction using AlphaFold2^46^, and subsequently modeled PEP into the allosteric pocket of each model by structural alignment with the previously solved mutant *Geobacillus stearothermophilus* PFK-1-PEP (PDB: 4I4I) complex (Figure 5A) (all-atom RMSD of the binding pocket = 0.371 Å; global RMSD = 0.493 Å).^41^ We optimized each ortholog’s PFK-1 model to a stable, energetically favorable conformation without steric clashes in the pocket using the relax application in the Rosetta macromolecular modeling software.^47^ Finally, to confirm that the models were reasonable, we calculated the ligand energy, RMSD, and TM-score of each model using US-align with the PEP-bound PFK-1 from *Geobacillus* as the reference structure and filtered out outliers or models with positive ligand energies (Figure 5B).^14,41,48^ Additionally, models were excluded if the ligand was observed to bind outside of the binding pocket after relaxing the structure. The final pool of PEP-bound bacterial PFK-1 structural models included 72 PFK-1 orthologs from 7 phyla and 39 orders. We calculated an average 50.2% sequence identity between any two PFK-1 from one phylum, vs. 29.54% between PFK-1 from different phyla, which suggested that our models sufficiently represented bacterial sequence diversity. ADP-bound PFK-1 models were prepared using the same strategy, using the crystal structure of ADP-bound *Escherichia coli* PFK-1 (PDB: 1PFK) as a reference for structural alignment. This resulted in a pool of 65 ADP-bound bacterial PFK-1 models from 4 phyla and 15 orders.

MAGPIE visualization of the PEP-bound PFK-1 models revealed four different PEP conformers among the 72 structures after binning by 0.4 Å global RMSD using our helper script. The top two conformers exhibited nearly identical spatial arrangement of hydrogen bond partners (Figures 6A, S5). Using a 1.5 Å threshold for distance among hotspot AAs and a minimum number of 15 AAs to form a hotspot, MAGPIE identifies two hotspots for each of the PEP conformers. Both hotspots for the second conformer pool are localized to the same region, which overlaps with one of the hotspots from the first conformer pool, relative to the ligands’ functional groups. However, MAGPIE found a unique hotspot for Conformer 1 near the phosphate group of PEP. These findings suggest that PEP can bind in different conformations that interact with spatially and biochemically distinct sets of AAs enriched in the allosteric site of PFK-1 orthologs, while also maintaining some conformer-independent trends. We hypothesize that the enrichment of Conformer 1 in our dataset is due to its ability to form stabilizing interactions with binding pocket residues in Hotspot 2 that are not possible for other PEP conformers.

By contrast, MAGPIE visualization of the ADP-bound PFK-1 models revealed four different ADP conformers among the 65 input structures after binning conformer pools using a 0.7 Å global RMSD for the ADP molecule (Figure 6B). Most models were binned into the first conformer category (n=58), so we focused our analysis on this ADP conformer. Using a 1.5 Å threshold for distance among hotspot AAs and a minimum number of 20 AAs to form a hotspot, MAGPIE identifies 6 hotspots in tight clusters around ADP. These hotspots localize around each functional group of the ADP molecule, with Hotspot 1 near the beta-phosphate, Hotspots 4 and 5 near the adenosine moiety, and Hotspot 2 near the ribose group, for example. Interestingly, the hydrophobic region that dominates ADP Hotspot 1 overlaps spatially with PEP Conformer 1’s unique hotspot. Differences in the PEP and ADP hotspots can be used to inform the rational design of small molecule binders to inhibit or activate bacterial PFK-1 orthologs using a similar mechanism as the enzyme’s natural effectors to artificially modulate the enzyme’s activity.

Further, using MAGPIE to compare PEP interactions in structural models of bacterial vs. metazoan PFK-1 orthologs could guide the design of bacteria-specific inhibitors as antibiotics that do not disrupt human PFK-1 in tissues.^49^

## Discussion

As highly accurate protein structure prediction methods spur the development of computational approaches to model and design protein-ligand binding interactions, the need for versatile tools to analyze the biochemical details of structural complexes *en masse* grows. Methods to easily compare protein interactions with other biomolecules will aid in biological discovery and therapeutics development. MAGPIE provides a practical way to explore the diverse interactions that natural and designed proteins alike can form with protein and small molecule ligands.

## Methods

### Data collection and curation

MAGPIE datasets of protein complex structures require at least one common target ligand. They can be computationally generated models (*e.g.*, homology models or designed complexes) or experimentally solved structures. Anyone can assemble a MAGPIE dataset using the PDB advanced search query builders. For the protein-protein complex case study, we used a 68-member subset of the previously compiled SARS-CoV-2 RBD-antibody dataset from Gowthaman *et al*.^34^ To assemble the small molecule ligand (COA) dataset included here, we used the chemical similarity search in the PDB with the PubChem identifier code for COA, 87642, as the chemical attribute. We defined the refinement resolution for the range 1.5-3 Å. The query resulted in >600 structural models, from which we randomly chose 199 models. More information about the models in this study is in section 8 of the supplementary material.

### Input structure standardization and alignment

Recognizing that input structures of protein-protein and protein-small molecule complexes for analysis by MAGPIE may include heterogeneous components, we prioritized flexibility in our structure preparation method (helper script MAGPIE_input_prep.py) by enabling a variety of user options to identify the target ligands and protein binders in the datasets. Users must define an input directory, an output directory, and identification options for both the target ligand and the protein binders. A pre-filter step can be applied to the PDBs entailed in a method called mesh search. Measured in Ångstroms, chain name(s), sequence fragment(s), ligand PDB code(s), and ligand indices can be supplied to conduct a nearest neighbor search. The input criteria become centers for a radial search in which anything outside an 8 Å radius will be filtered out. If any atom from another chain falls within the search radius, the structure to which it belongs is retained. This pre-filter step is used to narrow down undesired atoms that later filtering steps would fail to remove. For protein target ligands, users can define the chain name and/or all or part of the AA sequence as a string or file, optionally with a sequence identity percentage threshold. For small molecule target ligands, users can define the chain name and/or the PDB ID code and/or residue index. For the protein binders, regardless of target ligand type, users can define chain name(s) and/or protein sequence(s). If more than one identifier is provided for the target or binder, the user can also optionally define the order in which they are identified. All elements of the structure that are neither target ligand or protein binder are then deleted, and the protein complex chains are renamed such that all the target chains and ligand chains are respectively the same for all structural models in the set. If there is more than one chain defined as the binder or target, it is combined and renumbered as one chain. In the case of small molecule ligands, the atoms can also be renumbered and renamed to be consistent for all the models via a second-order connectivity graph using a bond length of 2.1 Å to find neighboring atoms. This method is recursively called to ensure atom identification and naming consistency.

Once the ligands and binders are standardized in the set of input structures, they are protonated and supplied to a structural alignment method that we implemented as a command line tool using the macromolecular modeling software PyMOL (Schrodinger) (helper script align_protein_chain.py for protein target ligands, and align_small_molecule.py for small molecule target ligands). For complexes that include protein target ligands, the input structures are returned after global structural alignment on the target. For small molecule ligands, there may be several ligand conformations represented in the dataset. Therefore, the user can define a maximum RMSD threshold (Å), and the script returns one or more directories, each containing sets of aligned structural models with target ligand conformations that fall within the RMSD threshold. Each cluster should be separately input to MAGPIE to visualize and analyze the local environment of each of its atoms.

### Performance, scalability, and web server implementation

MAGPIE is implemented in Python and uses multiple CPU processors to optimize speed. MAGPIE can be run on the cloud-based Google CoLab server if the input PDB dataset includes fewer than about 1000 structures. For private and/or larger datasets where limited computing resource allocations are a concern, we advise instead downloading the MAGPIE source code and running it locally and/or using the multithreaded implementation.

### Software accessibility

MAGPIE is available open source from the Glasgow Lab GitHub at https://github.com/glasgowlab/MAGPIE/tree/local-version and freely available to download with documentation and two dataset/usage examples. Local installation requirements are listed in section 9 of the supplementary material. A Google CoLab server running MAGPIE, where users can run the case studies or their own uploaded datasets, is also available at https://colab.research.google.com/github/glasgowlab/MAGPIE/blob/GoogleColab/MAGPIE_COLAB.ipynb. We provide helper scripts (MAGPIE_input_prep.py, align_protein_chain.py, and align_small_molecule.py) to clean, standardize, and align input PDB files for seamless usage in MAGPIE as described above, with documentation in section 6 of the supplementary material.

## Supplementary material description

Supplementary figures are available in section 1 of the supplementary material. A detailed description of the MAGPIE method, recommendations for preparing datasets for MAGPIE manually or with the provided tools, a list of additional features, a troubleshooting guide, detailed documentation for the helper scripts for preparing and cleaning input structures, information about multithreaded MAGPIE, and runtime benchmarks are provided in sections 2-7. Additional information about the structural models in the case studies is in section 8. Software dependencies and instructions for preparing a Conda environment to run MAGPIE are available in sections 9 and 10.

## Acknowledgements

The authors thank the Glasgow Lab and Dr. James Lucas for beta testing, feature requests, and bug reports.

## Abbreviations

MAGPIE: Mapping Areas of Genetic Parsimony In Epitopes
AA: amino acid
HA: heavy atom
MSA: multiple sequence alignment
RMSD: root-mean-square deviation
RBD: receptor binding domain
CDR: complementarity-determining region
COA: coenzyme A
PFK-1: phosphofructokinase-1
PEP: phosphoenolpyruvate
ADP: adenosine diphosphate

## References

1. Crooks GE, Hon G, Chandonia J-M, Brenner SE (2004) WebLogo: A Sequence Logo Generator. Genome Res. 14:1188–1190.

2. Larkin MA, Blackshields G, Brown NP, Chenna R, McGettigan PA, McWilliam H, Valentin F, Wallace IM, Wilm A, Lopez R, et al. (2007) Clustal W and Clustal X version 2.0. Bioinformatics 23:2947–2948.

3. Sievers F, Wilm A, Dineen D, Gibson TJ, Karplus K, Li W, Lopez R, McWilliam H, Remmert M, Söding J, et al. (2011) Fast, scalable generation of high-quality protein multiple sequence alignments using Clustal Omega. Molecular Systems Biology 7:539.

4. Katoh K, Standley DM (2013) MAFFT Multiple Sequence Alignment Software Version 7: Improvements in Performance and Usability. Molecular Biology and Evolution 30:772–780.

5. Wallace IM, O’Sullivan O, Higgins DG, Notredame C (2006) M-Coffee: combining multiple sequence alignment methods with T-Coffee. Nucleic Acids Research 34:1692–1699.

6. Brown NP, Leroy C, Sander C (1998) MView: a web-compatible database search or multiple alignment viewer. Bioinformatics 14:380–381.

7. Hagopian R, Davidson JR, Datta RS, Samad B, Jarvis GR, Slander K (2010) SATCHMO-JS: a webserver for simultaneous protein multiple sequence alignment and phylogenetic tree construction. Nucleic Acids Research 38:W29–W34.

8. Brown BP, Mendenhall J, Meiler J (2019) BCL::MolAlign: Three-Dimensional Small Molecule Alignment for Pharmacophore Mapping. J. Chem. Inf. Model. 59:689–701.

9. Roy A, Skolnick J (2015) LIGSIFT: an open-source tool for ligand structural alignment and virtual screening. Bioinformatics 31:539–544.

10. McGaughey GB, Sheridan RP, Bayly CI, Culberson JC, Kreatsoulas C, Lindsley S, Maiorov V, Truchon J-F, Cornell WD (2007) Comparison of Topological, Shape, and Docking Methods in Virtual Screening. J. Chem. Inf. Model. 47:1504–1519.

11. Lemmen C, Lengauer T, Klebe G (1998) FlexS: A Method for Fast Flexible Ligand Superposition. J. Med. Chem. 41:4502–4520.

12. Hawkins PCD, Skillman AG, Nicholls A (2007) Comparison of Shape-Matching and Docking as Virtual Screening Tools. J. Med. Chem. 50:74–82.

13. Hönig SMN, Lemmen C, Rarey M (2023) Small molecule superposition: A comprehensive overview on pose scoring of the latest methods. WIREs Computational Molecular Science 13:e1640.

14. Zhang Y, Skolnick J (2005) TM-align: a protein structure alignment algorithm based on the TM-score. Nucleic Acids Research 33:2302–2309.

15. Holm L Using Dali for Protein Structure Comparison. In: Gáspári Z, editor. Structural Bioinformatics: Methods and Protocols. Methods in Molecular Biology. New York, NY: Springer US; 2020. pp. 29–42. Available from: 10.1007/978-1-0716-0270-6_3

16. van Kempen M, Kim SS, Tumescheit C, Mirdita M, Lee J, Gilchrist CLM, Söding J, Steinegger M (2023) Fast and accurate protein structure search with Foldseek. Nat Biotechnol:1–4.

17. Chen EA, Porter LL (2023) SSDraw: Software for generating comparative protein secondary structure diagrams. Protein Science 32:e4836.

18. Evans R, O’Neill M, Pritzel A, Antropova N, Senior A, Green T, Žídek A, Bates R, Blackwell S, Yim J, et al. (2022) Protein complex prediction with AlphaFold-Multimer.: 2021.10.04.463034. Available from: https://www.biorxiv.org/content/10.1101/2021.10.04.463034v2

19. Bryant P, Pozzati G, Elofsson A (2022) Improved prediction of protein-protein interactions using AlphaFold2. Nat Commun 13:1265.

20. Zhang QC, Petrey D, Deng L, Qiang L, Shi Y, Thu CA, Bisikirska B, Lefebvre C, Accili D, Hunter T, et al. (2012) Structure-based prediction of protein–protein interactions on a genome-wide scale. Nature 490:556–560.

21. Hwang H, Dey F, Petrey D, Honig B (2017) Structure-based prediction of ligand–protein interactions on a genome-wide scale. Proceedings of the National Academy of Sciences 114:13685–13690.

22. Trudeau SJ, Hwang H, Mathur D, Begum K, Petrey D, Murray D, Honig B (2023) PrePCI: A structure-and chemical similarity-informed database of predicted protein compound interactions. Protein Science 32:e4594.

23. Petrey D, Zhao H, Trudeau SJ, Murray D, Honig B (2023) PrePPI: A Structure Informed Proteome-wide Database of Protein–Protein Interactions. Journal of Molecular Biology 435:168052.

24. Kynast JP, Höcker B (2023) Atligator Web: A Graphical User Interface for Analysis and Design of Protein–Peptide Interactions. BioDesign Research 5:0011.

25. Ferruz N, Schmidt S, Höcker B (2021) ProteinTools: a toolkit to analyze protein structures. Nucleic Acids Res 49:W559–W566.

26. Bentley JL (1975) Multidimensional binary search trees used for associative searching. Commun. ACM 18:509–517.

27. Sievert C (2020) Interactive Web-Based Data Visualization with R, plotly, and shiny. Available from: https://plotly-r.com

28. Sayle R (1995) Rasmol.

29. Tareen A, Kinney JB (2020) Logomaker: beautiful sequence logos in Python. Bioinformatics 36:2272–2274.

30. Pedregosa F, Varoquaux G, Gramfort A, Michel V, Thirion B, Grisel O, Blondel M, Prettenhofer P, Weiss R, Dubourg V, et al. (2011) Scikit-learn: Machine Learning in Python. Journal of Machine Learning Research 12:2825–2830.

31. Ester M, Kriegel H-P, Sander J, Xu X A density-based algorithm for discovering clusters in large spatial databases with noise. In: Proceedings of the Second International Conference on Knowledge Discovery and Data Mining. KDD’96. Portland, Oregon: AAAI Press; 1996. pp. 226–231.

32. Lisanza SL, Gershon JM, Tipps S, Arnoldt L, Hendel S, Sims JN, Li X, Baker D Joint Generation of Protein Sequence and Structure with RoseTTAFold Sequence Space Diffusion. Biochemistry; 2023. Available from: http://biorxiv.org/lookup/doi/10.1101/2023.05.08.539766

33. Watson JL, Juergens D, Bennett NR, Trippe BL, Yim J, Eisenach HE, Ahern W, Borst AJ, Ragotte RJ, Milles LF, et al. (2023) De novo design of protein structure and function with RFdiffusion. Nature 620:1089–1100.

34. Gowthaman R, Guest JD, Yin R, Adolf-Bryfogle J, Schief WR, Pierce BG (2021) CoV3D: a database of high resolution coronavirus protein structures. Nucleic Acids Research 49:D282– D287.

35. Glasgow A, Glasgow J, Limonta D, Solomon P, Lui I, Zhang Y, Nix MA, Rettko NJ, Zha S, Yamin R, et al. (2020) Engineered ACE2 receptor traps potently neutralize SARS-CoV-2. Proceedings of the National Academy of Sciences 117:28046–28055.

36. Remesh SG, Merz GE, Brilot AF, Chio US, Rizo AN, Pospiech TH, Lui I, Laurie MT, Glasgow J, Le CQ, et al. (2023) Computational pipeline provides mechanistic understanding of Omicron variant of concern neutralizing engineered ACE2 receptor traps. Structure 31:253–264.e6.

37. Kariolis MS, Miao YR, Jones DS, Kapur S, Mathews II, Giaccia AJ, Cochran JR (2014) An engineered Axl “decoy receptor” effectively silences the Gas6-Axl signaling axis. Nat. Chem. Biol. 10:977–983.

38. Chan KK, Dorosky D, Sharma P, Abbasi SA, Dye JM, Kranz DM, Herbert AS, Procko E (2020) Engineering human ACE2 to optimize binding to the spike protein of SARS coronavirus 2. Science 369:1261–1265.

39. Dauparas J, Lee GR, Pecoraro R, An L, Anishchenko I, Glasscock C, Baker D (2023) Atomic context-conditioned protein sequence design using LigandMPNN.: 2023.12.22.573103. Available from: https://www.biorxiv.org/content/10.1101/2023.12.22.573103v1

40. Lucas JE, Kortemme T (2020) New computational protein design methods for de novo small molecule binding sites. PLOS Computational Biology 16:e1008178.

41. Mosser R, Reddy MCM, Bruning JB, Sacchettini JC, Reinhart GD (2013) Redefining the Role of the Quaternary Shift in Bacillus stearothermophilus Phosphofructokinase. Biochemistry 52:5421–5429.

42. Shirakihara Y, Evans PR (1988) Crystal structure of the complex of phosphofructokinase from Escherichia coli with its reaction products. Journal of Molecular Biology 204:973–994.

43. Tian T, Wang C, Wu M, Zhang X, Zang J (2018) Structural Insights into the Regulation of *Staphylococcus aureus* Phosphofructokinase by Tetramer–Dimer Conversion. Biochemistry 57:4252–4262.

44. Paricharttanakul NM, Ye S, Menefee AL, Javid-Majd F, Sacchettini JC, Reinhart GD (2005) Kinetic and Structural Characterization of Phosphofructokinase from Lactobacillus bulgaricus. Biochemistry 44:15280–15286.

45. Currie MA, Merino F, Skarina T, Wong AHY, Singer A, Brown G, Savchenko A, Caniuguir A, Guixé V, Yakunin AF, et al. (2009) ADP-dependent 6-Phosphofructokinase from *Pyrococcus horikoshii* OT3. Journal of Biological Chemistry 284:22664–22671.

46. Jumper J, Evans R, Pritzel A, Green T, Figurnov M, Ronneberger O, Tunyasuvunakool K, Bates R, Žídek A, Potapenko A, et al. (2021) Highly accurate protein structure prediction with AlphaFold. Nature 596:583–589.

47. Alford RF, Leaver-Fay A, Jeliazkov JR, O’Meara MJ, DiMaio FP, Park H, Shapovalov MV, Renfrew PD, Mulligan VK, Kappel K, et al. (2017) The Rosetta All-Atom Energy Function for Macromolecular Modeling and Design. J. Chem. Theory Comput. 13:3031–3048.

48. Zhang C, Shine M, Pyle AM, Zhang Y (2022) US-align: universal structure alignments of proteins, nucleic acids, and macromolecular complexes. Nat Methods 19:1109–1115.

49. McNae IW, Kinkead J, Malik D, Yen L-H, Walker MK, Swain C, Webster SP, Gray N, Fernandes PM, Myburgh E, et al. (2021) Fast acting allosteric phosphofructokinase inhibitors block trypanosome glycolysis and cure acute African trypanosomiasis in mice. Nat Commun 12:1052.

