## Supplemental information for "MAGPIE: an interactive tool for visualizing and analyzing protein-ligand interactions"

### 1. Supplemental figures

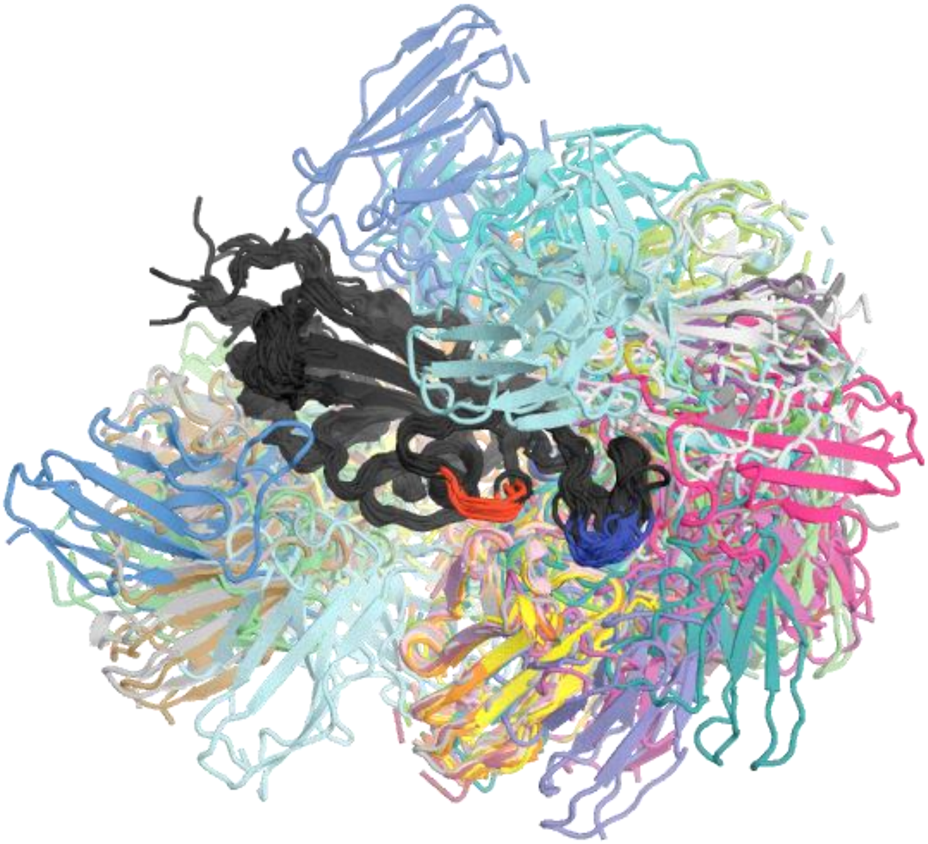

**Figure S1. Structural models of all the antibody-SARS-CoV-2 spike receptor binding domain (RBD) complexes in our study.** The models are aligned on the RBD (colored black). RBD Loop 1 is colored red and Loop 2 is colored blue.

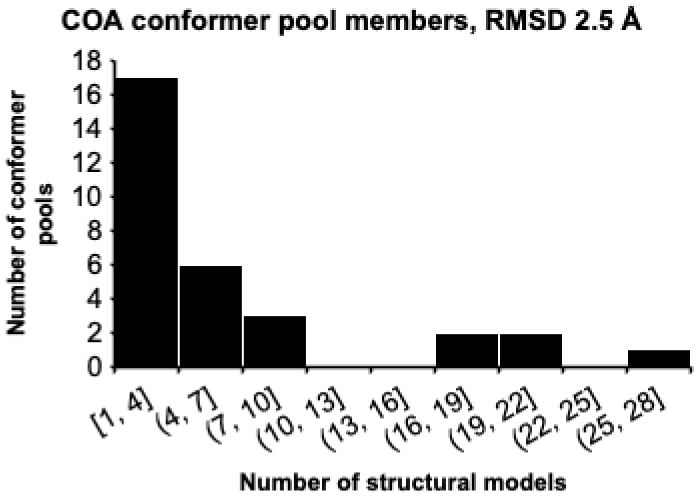

**Figure S2. COA conformer pools.** The 199 COA structural models were aligned using our helper script small_molecule_align.py and separated into 31 conformer pools using an RMSD threshold of 2.5 Å. Conformer pools ranged in size from just one model to 27 models. Seven conformer pools contained a single model with unique COA conformers, while 5 conformer pools contained 17 or more models.

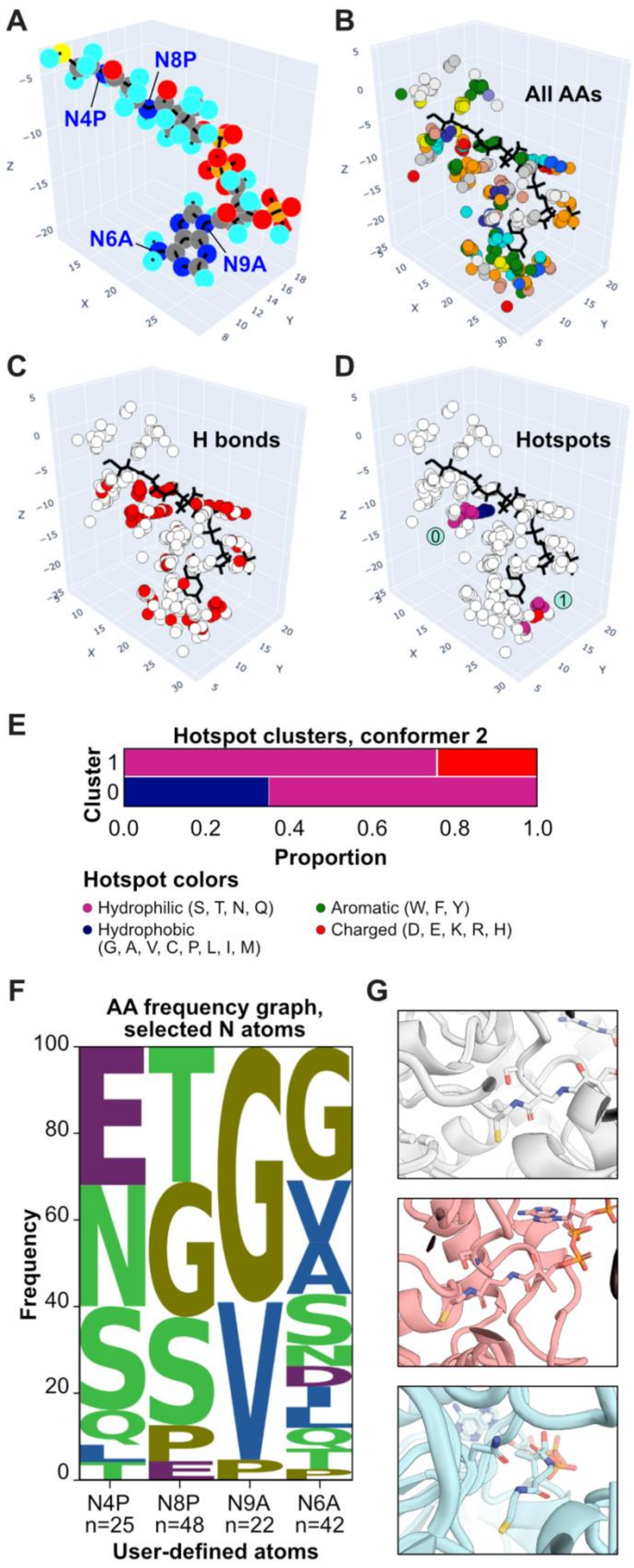

**Figure S3. MAGPIE analysis of COA Conformer Pool 2. (A)** MAGPIE-generated structural representation of Conformer 2. **(B)** The 21 models in Conformer Pool 2 within 5 Å of the COA conformer, colored by the Amino color code, as in Figure 2. **(C)** Hydrogen bond partners in red. **(D, E)** Hotspots and cluster analysis, as defined by DBSCAN settings eps = 2, min_samples = 20. Two hotspots were found, with different AA compositions as compared to COA Conformer 1. **(F)** AA frequency graph showing the local environment for four nitrogen atoms in COA Conformer Pool 2. **(G)** Examples of structural models from the conformer pool, zoomed in on interaction partners near COA nitrogen atom N4P.

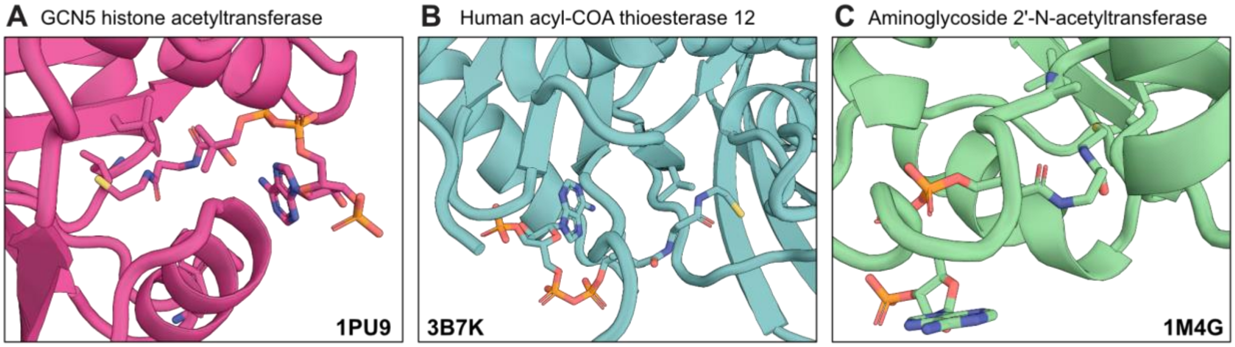

**Figure S4. Conformer Pool 1 structural models.** Examples of specific enzymes from the pool showing the proximity of the N4P HA to hydrophobic AAs. The PDB IDs are in the corners of each panel.

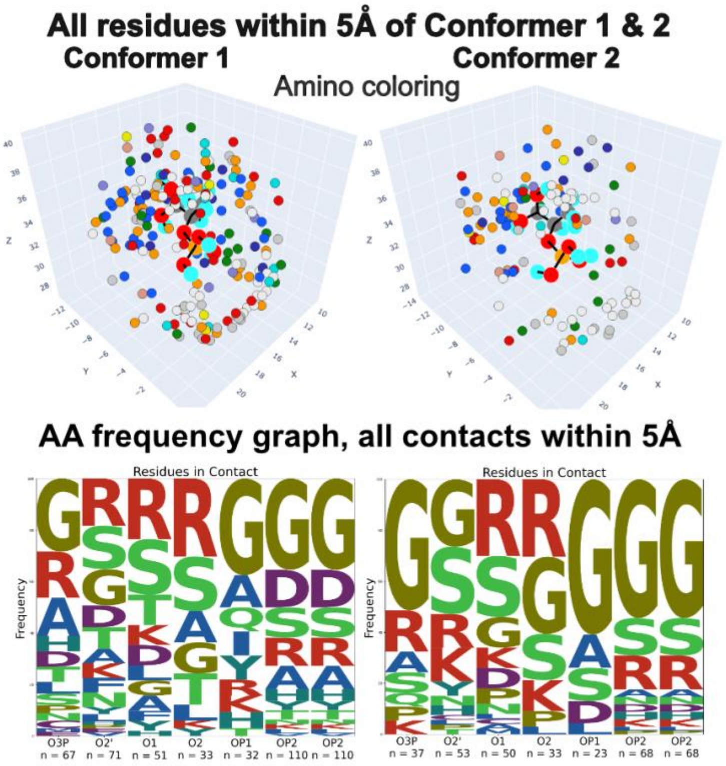

**Figure S5. Local environment for PEP Conformers 1 and 2.** MAGPIE 3D visualization of AAs within 8 Å of PEP conformer 1 and 2 and corresponding AA frequency graphs.

### 2. MAGPIE: detailed methods

**Determining residues in contact with target ligand positions**

Protein Data Bank (PDB) files are parsed using the Biopython PDB module.[1] MAGPIE extracts the location of all alpha carbons from the user-provided chain indices. By using a K-DTree neighbor search with the given distance, MAGPIE stores residue information, including all atomic coordinates, found to be in contact and defined by the distance between two alpha carbon atoms to be less than or equal to the given distance. Only residues from the target chain that are in contact with at least one residue from the binder chain are stored, and vice versa. This avoids unnecessary calculations in future steps. MAGPIE performs a similar process for small molecule ligands using each atom as a residue and following the described pipeline.

**Determining hydrogen bonds (H bonds)**

Atoms are classified either as H bond acceptors or donors based on the following criteria: any H atom attached to an electronegative atom (O, N) was defined as a donor, and each atom defined as an electronegative atom was defined as an acceptor. These pairs were compared across all residues in contact with the target ligand. To determine the presence of H bonds, MAGPIE calculates two vectors defined as:

$$A\vec{D}_1 = A - D_1$$

$$D_2\vec{D}_1 = D_2 - D_1$$

where A is the acceptor coordinates, D1 is the H atom donor coordinates, and D2 is the respective electronegative atom donor coordinates. These vectors are centered around D1 and therefore the angle between them can be used to calculate the angle center at D1.

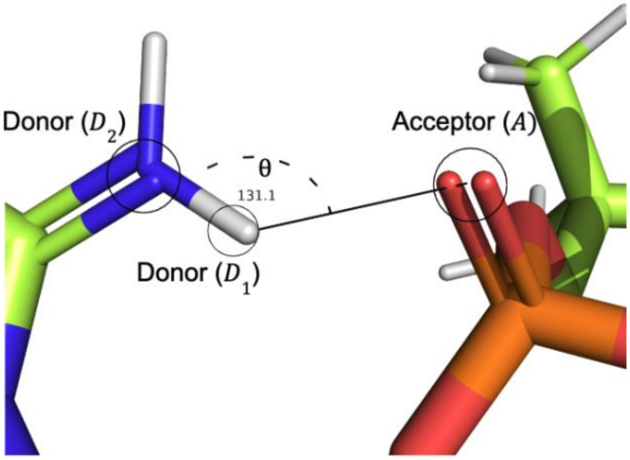

$$\theta = \arccos\left(\frac{\vec{AD_1} \cdot \vec{D_2D_1}}{\|\vec{AD_1}\|\|\vec{D_2D_1}\|}\right) \quad (1)$$

An H bond is defined by bond angle $120° \leq \theta \leq 240°$ and bond length $2\ \text{Å} \leq AD_1 \leq 4\ \text{Å}$.[2]

**Determining salt bridges**

Salt bridges are defined by the distance between a positively charged atom coordinates $SB_+$ and a negatively charged atom coordinates $SB_-$ where:

$$\vec{SB} = \|SB_+ - SB_-\| \quad (2)$$

Every negatively charged atom is compared to every other possible charged atom, as shown in Table 1.

| Charge | Amino Acid | Atom IDs |
|---|---|---|
| +1 | LYS | NZ |
| +1 | ARG | NH1, NH2, NE |
| +1 | HIS | ND1, NE2 |
| -1 | GLU | OE1, OE2 |
| -1 | ASP | OD1, OD2 |
| -1 | CYS | SG |
| -1 | TYR | OH |

Table 1: Amino Acids and Their Charged Atoms

A salt bridge is defined by $1\ \text{Å} \leq SB \leq 4.5\ \text{Å}$. Salt bridges are only calculated for protein-protein interactions.[3]

**Determining the percentage of AA neighbors per position**

To calculate the AA frequency graph, MAGPIE first defines a reference structure to use. For protein-protein interactions, it uses chain length (how many AAs with a given chain index) to obtain the chain with the largest chain length across all PDB inputs. MAGPIE helper scripts make sure that small molecule ligands have the same number of atoms, and atom IDs, therefore MAGPIE chooses an arbitrary reference.

MAGPIE then runs a K-DTree neighbor search with the given distance to find residues that are in contact with the input residue or atom. It calculates frequencies by dividing the number of each AA type by the total number of AAs found.

### 3. Recommendations for preparing structural models for MAGPIE

##### Compiling models

Any Protein Data Bank (PDB)-format protein complex structures that include at least one protein can be used as input data for MAGPIE. They can be experimentally solved or computationally generated structural models. We have used MAGPIE to analyze and visualize computationally designed protein complex models, experimentally solved protein complexes from the Protein Data Bank[4], computationally predicted structural models of protein oligomers[5], and docked ligand-bound proteins[6]. MAGPIE can process datasets with thousands of models. The likelihood that MAGPIE detects hotspots will scale with the number of structures, so the user should adjust the hotspot parameters accordingly (see also section 4, Hotspot identification and visualization).

##### Cleaning models

All models supplied to MAGPIE should include the target ligands and protein binders on separate chains. The chain label for the target ligand and the protein binders should be different, but the chain label for the target ligand should be the same for all models, and the chain label for the protein binders should be the same for all models. MAGPIE does not require the protein binders to have the same number of amino acids, but the target ligand should ideally have identical amino acid or atom numbering. MAGPIE will run with partial ligands or missing amino acids, but the user should check the input structures before interpreting the results. Other chains that the user does not specify as the target ligand or protein binder are permitted but will be ignored by MAGPIE.

Three optional helper scripts for preparing PDB input files for MAGPIE are provided on GitHub (https://github.com/glasgowlab/MAGPIE):

1. the cleaning script, MAGPIE_input_prep.py
2. an alignment script for structural models with target ligands that are proteins, align_protein_chain.py
3. the alignment script for structural models with target ligands that are small molecules, align_small_molecule.py

See the GitHub README file or section 6 of this document for the full documentation and instructions for using the helper scripts to prepare input structures for MAGPIE.

##### Structural alignment

MAGPIE requires that the structures in the dataset are aligned on the target ligand. Any macromolecular modeling software can be used to align the structures: for example, the “align” or “cealign” commands can be used in PyMOL[7], or the alignment commands in ChimeraX[8]. (The provided alignment scripts use the PyMOL API in the background.)

**Number of input models**

MAGPIE is compatible with any number of inputs, but the multithreaded local version of the software has improved performance for >1000 input PDB files as compared to the Google CoLab implementation. Both versions are available as branches on GitHub (https://github.com/glasgowlab/MAGPIE). See section 7 of this document for information about running multithreaded MAGPIE and for runtime benchmarks.

### 4. Additional features

##### Hotspot identification and visualization

MAGPIE uses the Python scikit-learn implementation of DBSCAN[9,10] to find hotspot clusters, with two settings.

- *eps* is the maximum distance between two AAs for either to be considered in a cluster with the other. Default value, 2 Å.
- *min_samples* is the number of AA in a cluster to define a hotspot. Default value, 15.

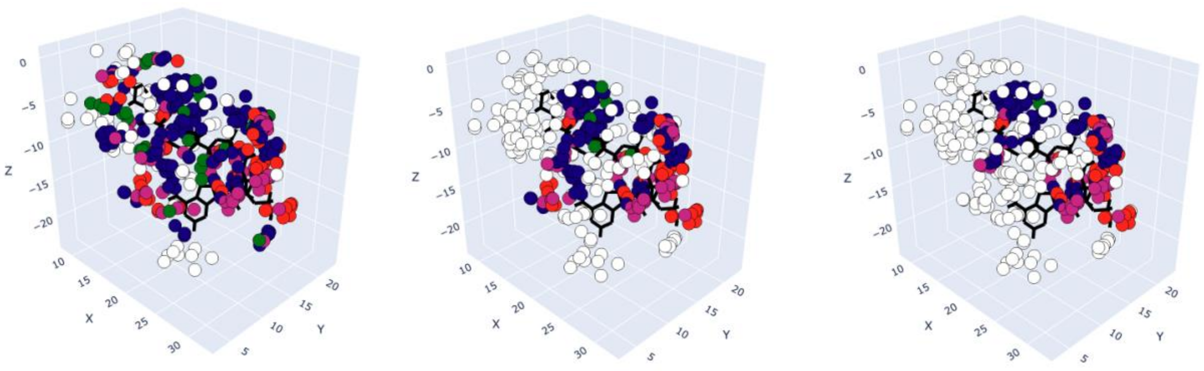

With all residues within 5 Å of the target ligand (black sticks) and eps = 2,

Left: min_samples = 5 → **19 hotspots**

Middle: min_samples = 10 → **12 hotspots**

Right: min_samples = 25 → **6 hotspots**

##### Metadata export

MAGPIE can produce two exported .csv files for an organized, detailed summary table of the interactions found in each dataset.

Hotspot metadata: this file contains each hotspot residue and index and tabulates the number of each AA in each hotspot, the fractional contribution of each AA to each hotspot, and the total number of AAs in each hotspot.

Polar and proximity metadata: for each AA that MAGPIE finds to interact with the target ligand, this file contains:

- X, Y, Z coordinates of its Cα atom
- chain name, residue index, and residue name
- HEX color codes for the Shapely and Amino color schemes

- file name for the originating structural model (.pdb)
- backbone and side chain H bond information - in both cases, only atoms corresponding to donors and acceptors from H bonds defined by Equation (1) are stored. The distinction between backbone H bond and side chain H bond is defined by the atom originating from the target chain.
- atoms involved in the formation of salt bridges as defined by Equation (2).
- the residue index for the AA in the target ligand that it interacts with, in the originating structural model
- hotspot cluster index (-1 = not in a hotspot)
- AA class (hydrophobic, aromatic, charged, hydrophilic)

**Toggle chains**

MAGPIE can visualize the target ligand or the binding partners alone.

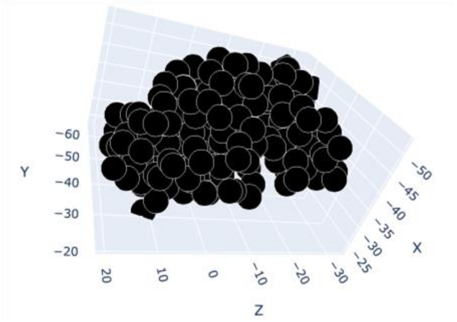

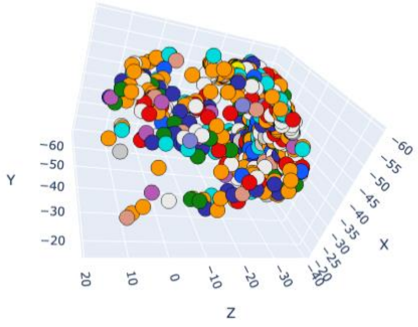

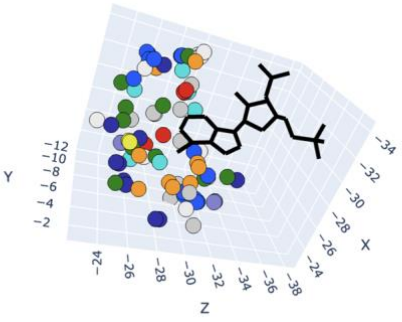

**Protein binder AA color scheme**

MAGPIE can visualize amino acids in the protein binding partners using the “Amino” or “Shapely” color schemes.

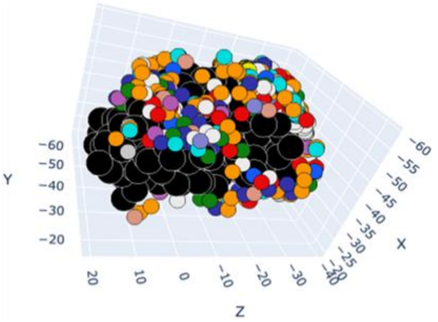

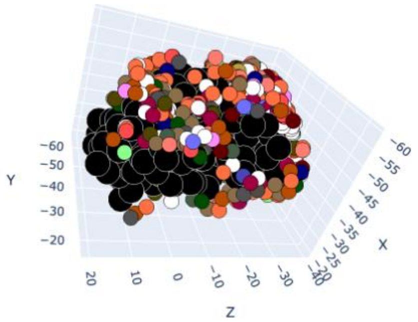

### 5. Troubleshooting guide

| | | |
|---|---|---|
| **Loading and processing files** | IndexError: single positional indexer is out-of-bounds | <u>Possible fixes:</u><br>Check that .zip files are not protected or corrupted.<br><br>Check that the files are updated to the right directory:<br>- For CoLab, check that the zip file is located in the 'temp' folder.<br>- For CoLab: only run Step 1 once.<br>- For local: confirm that the right directory is inputted.<br><br>Step 2 prints out a list with the location of the path. Make sure you can see your files.<br>- If the files do not update, you can move them from the Google Drive content folder.<br>- Refresh folder if not unzipping correctly.<br>Check for a print statement confirming that the files were successfully inputted. |
| **Plotting 3D visualization** | Exception: Protein Example contains chain A, which is not defined, please remove or rename chain.<br><br>ValueError: x must consist of vectors of length 3 but has shape (0,)<br><br>Unable to solve small-molecule ligand bonds. | <u>Possible fixes:</u><br>This occurs when the program cannot find any entries corresponding to the input chains or residues.<br>- Check to confirm the correct characters, capitalization, and no spaces.<br>- Check that you only use the small molecule ligand option if you are plotting a ligand. |
| **Plotting MAGPIE AA frequency graphs** | TypeError: ufunc 'isfinite' not supported for the input types, and the inputs could not be safely coerced to any supported types according to the casting rule ''safe'' | <u>Possible fixes:</u><br>Three possible reasons:<br>- There are no residues in close proximity.<br>- The inputs are wrong, so check if they are properly separated by commas and that the names are correct.<br>- Check for extra spaces. |
| **Wrong version of MAGPIE** | UsageError: Line magic function '%%capture' not found. | <u>Possible fixes:</u><br>Make sure you are using the correct version of MAGPIE. MAGPIE collab will not execute properly in a local jupyter notebook setting. Clone the appropriate repository. |

### 6. Helper scripts to prepare input files for MAGPIE

Recognizing that protein structure files downloaded from the PDB or prepared using custom programs will have many different formats, we provide the helper Python script **MAGPIE_input_prep.py** in the Github GoogleCoLab branch to help users prepare input PDB files for MAGPIE. The helper script takes an input PDB file or directory of PDB files, an output directory, and identifies information about the protein binders and the target ligands. It outputs reformatted, renumbered PDB files in which the protein binder is on one chain and the target ligand is on another chain. The output files are found in the user-specified output directory with the suffix "_cleaned.pdb".

**Usage**

```
MAGPIE_input_prep.py [-h] [-S BINDER_SEQS] [-s TARGET_PROTEIN_SEQS] [-
F BINDER_SEQ_FA] [-f TARGET_PROTEIN_SEQ_FA] [-L TARGET_SM_3_NAMES] [-I
TARGET_SM_INDEX] [-C BINDER_CHAINS] [-c TARGET_PROTEIN_CHAINS][-l
TARGET_SM_CHAINS] [-U {chains,sequences}] [-u {chains,name}] [-d
SEQ_IDENTITY] -i INPUT_PATH -o OUTPUT_PATH [-N
TARGET_PROTEIN_CHAIN_RENAME] [-n TARGET_SM_CHAIN_RENAME] [-b
BINDER_CHAIN_RENAME] [-t TAKE_FIRST_SM_ONLY] [-T NAME_SM_ATOMS_SAME]
[-r SM_LIGAND_REFERENCE_PATH] [-B BOND_LENGTH] [-m SEARCH_RADIUS] [-M
MESH_SEARCH] [-A SEQ_TARGET_ALIGN] [-a SEQ_TARGET_REF_PDB]
```

**Argument definitions**

1. `-h, --help`

   Show this help message and exit.

2. `-S BINDER_SEQS, --binder_seqs BINDER_SEQS`

   The sequence(s) of the binder used for identification.

3. `-s TARGET_PROTEIN_SEQS, --target_protein_seqs TARGET_PROTEIN_SEQS`

   The sequence(s) of the target protein used for identification.

4. `-F BINDER_SEQ_FA, --binder_seq_fa BINDER_SEQ_FA`

   The sequence file (fasta format) of the binder used for identification.

5. `-f TARGET_PROTEIN_SEQ_FA, --target_protein_seq_fa TARGET_PROTEIN_SEQ_FA`

   The sequence file (fasta format) of the target protein used for identification.

6. `-L TARGET_SM_3_NAMES, --target_sm_3_names TARGET_SM_3_NAMES`

The three letter code of the small molecule(s).

7. `-I TARGET_SM_INDEX, --target_sm_index TARGET_SM_INDEX`

   The index of the small molecule(s). Must be used in tandem with chain or name.

8. `-C BINDER_CHAINS, --binder_chains BINDER_CHAINS`

   The chain(s) of the binder used for identification.

9. `-c TARGET_PROTEIN_CHAINS, --target_protein_chains TARGET_PROTEIN_CHAINS`

   The chains of the target protein used for identification.

10. `-l TARGET_SM_CHAINS, --target_sm_chains TARGET_SM_CHAINS`

    The chain(s) of the small molecule used for identification.

11. `-U {chains,sequences}, --search_first_protein {chains,sequences}`

    If using both chains and sequences search, what should be used to be filtered first? Choices: "chains" or "sequences" Default: "chains"

12. `-u {chains,name}, --search_first_sm {chains,name}`

    If using both chains and sequences search, what should be used to be filtered first? Choices: "chains" or "sequences" Default: "chains"

13. `-n TARGET_SM_CHAIN_RENAME, --target_sm_chain_rename TARGET_SM_CHAIN_RENAME`

    What the target small molecule output chain should be named. Default: "B"

14. `-N TARGET_PROTEIN_CHAIN_RENAME, --target_protein_chain_rename TARGET_SM_CHAIN_RENAME`

    What the target protein output chain should be named. Default: "A"

15. `-b BINDER_CHAIN_RENAME, --binder_chain_rename BINDER_CHAIN_RENAME`

    What the binder output chain should be named. Default: "C"

16. `-t TAKE_FIRST_SM_ONLY, --take_first_sm_only TAKE_FIRST_SM_ONLY`

    Should we only take the first instance of the ligand?

17. `-T NAME_SM_ATOMS_SAME, --name_sm_atoms_same NAME_SM_ATOMS_SAME`

    Should we rename all matching ligands with the same atom names? Uses the first file as a reference.

18. `-r SM_LIGAND_REFERENCE_PATH, --sm_ligand_reference_path SM_LIGAND_REFERENCE_PATH`

    Path of the reference file for renaming the small molecule ligands. Default is the first file found with Python's list directory function.

19. `-B BOND_LENGTH, --bond_length BOND_LENGTH`

    Distance that defines a bond between 2 atoms for chemical graphs. Default: 2.1

20. `-m SEARCH_RADIUS, --search_radius SEARCH_RADIUS`

    Distance that is considered for finding neighboring atoms in the mesh search. Default: 8

21. `-M MESH_SEARCH, --mesh_search MESH_SEARCH`

    The chains, sequence(s), small molecule name(s), small molecule index(es) for the mesh filter. Example: 'A,B;AWTRWARE,AWAWAWAW;TPA,ATP;1,2'

22. `-A SEQ_TARGET_ALIGN, --seq_target_align SEQ_TARGET_ALIGN`

    Should we align the target protein in sequence space? Results in PDB numbering via alignment. Do not use it for small molecule ligands. Choices: "1v1", "MSA"

23. `-a SEQ_TARGET_REF_PDB, --seq_target_ref_pdb SEQ_TARGET_REF_PDB`

    Reference structure for target protein in seq_target_align.

24. `-d SEQ_IDENTITY, --seq_identity SEQ_TARGET_REF_PDB`

    Sequence identity threshold for finding similar chains. Default: 95%

**Required arguments**

There are only two required command-line options. The first required option is the input, which is specified by the -i, --input_path. The input can either be a path to a single PDB file or to a directory holding one or more pdb files. The other required argument is the output which can be specified by the -o, --output_path option. The output must be a directory that has already been created.

**Mesh search**

Mesh search takes precedence over other search options. The user defines atoms that will become centers for a 3D radial search. The search is inclusive of the center points; any other atom found in the default radius of 8 Å will be considered part of the search results.

This search takes in two main command-line options:

1) -m, --search_radius

2) -M, --mesh_search

The --mesh_search option has four sub-options, all of which are separated by semicolons. An example input is the following: “A,B;AWTRWARE,AWAWAWAW;TPA,ATP;1,2”. The “A,B” in this search means we are looking for all atoms located in chain A and chain B to be considered part of the search centers. “AWTRWARE, AWAWAWAW” are sequences to be searched for to be considered as a center. The -d, --seq_identity option can be used for the percentage of sequence identity needed to be selected. If that sequence is found, the protein that it is a part of will be considered as part of the search center. “TPA,ATP” are small molecules that will be searched. If --take_first_sm_only is set to false, it will use all ligands it finds with the name provided by the user to be part of the search center; however, if it is not set, only the first instance of the molecule will be used as the center. The last part, “1,2” specifies specific PDB numbers used for small molecules. If the user had 4 TPA molecules, the ones with ID 1 and 2 would be selected. If they only want to search chains, the following input can be used: “A,B;;;”. The other blank arguments must have semicolons for this to work properly. The small molecule residue number must be used only if the small molecule names are given. The other three parameters can be used independently.

The --search_radius is defaulted to 8 Å. This option can be used to change the size of the search radius.

**Chain search**

The chain search allows the user to search for binders, small molecules, or target proteins on specific chains. The -C, --binder_chains option specifies what chains the binders are on, while the -c, --target_protein_chains option specifies the target, and -l, --target_sm_chains the small molecule. The chain search options can include multiple chains as long as you split the chain with a comma: “A,B”.

**Sequence search**

The sequence search allows the user to search for specific sequences in the PDB file. You can search for binders using the following arguments: -S, --binder_seqs for a few numbers of sequences: “AWTRWARE,AWAWAWAW”. Alternatively, you can use -F, --binder_seq_fa to give a path to a .fasta file for a lot of sequences. The equivalent target selection options are the following: -s, --target_protein_seq, and f, --target_protein_seq_fa. You can change the sequence identity needed to be considered a match by using the - d, --seq_identity option. If the user only specifies the target sequence and not binder options (chain or sequence), the program will default to all other protein atoms not containing the binder sequence. Also, by default, the program will use only the first instance of the sequence it finds and discard the rest of the proteins containing the sequence.

**Small molecule (SM) search**

The SM search allows the user to search for small molecules by name and index. The user can search for a target small molecule ligand by using the following option: -L, --target_sm_3_names, where multiple ligands can be specified by commas: “TPA,ATP”. The user can also include the index of the small molecules to further filter. The user must give a name for the -I, --target_sm_index option to work. An example input for the -I would be “1,2,5”. If the user does not turn off the -t, --take_first_sm_only option, only the first instance of the small molecule will be taken regardless of whether there are multiple small molecules or if the user picks specific indices.

**Order of searches**

Mesh search will take precedence over chain, sequence, and SM search. The mesh search will first pass over and filter the PDB file to only include atoms in your search radius and center points. Following the mesh search is either the chain search or sequence search for the binder and target protein. The order of the filter can be changed using the -U, --search_first_protein option. The order of the chain search and SM name search can also be determined by the -u, --search_first_sm option. By default, chains will be filtered before the sequence/SM name.

**Renaming the output chains**

If the user wants to rename the chain of the target, SM, and binder to something other than the default, A,B,C, they can use the -N, --target_protein_chain_rename for the target, n, --target_sm_chain_rename for the SM, and b, --binder_chain_rename for the binder.

**Renaming small molecule atoms**

If the user is filtering for small molecules, it may be helpful to rename the molecules to have all the same atom names for later steps. By default, the program will use the default atom names for the small molecules; however, if they should be named the same, the -T, --name_sm_atoms_same option can be used. If the option is selected, it will use the first PDB that has the target ligand as the reference PDB for the atom names of the small molecules. The user can define the reference PDB instead by using the -r, --sm_ligand_reference_path and providing a PDB path. The rename works by creating a second-order connectivity graph using a bond length of 2.1 Å to find neighboring atoms. This method is recursively called to create a connection of connections to distinguish the atoms. The length of the bond can be changed by using the -B, --bond_length option.

**MSA and 1V1 alignment**

If the user wants the target protein to be aligned either by a certain input structure's sequence or by a global MSA sequence, the -A, --seq_target_align can be applied. If they want an MSA for the alignment, they should ensure the argument is set to MSA. If they want to align to the longest sequence in a PDB compared to a reference, they can set the mode to 1v1. By default, the 1v1 mode will take the first PDB it finds as the reference; however, a reference structure/sequence can also be defined using the -a, --seq_target_ref_pdb. The MSA option will also output a sequence logo at the end as well as two additional files needed to run the Muscle MSA program.

**Examples**

1. The user has multiple PDB files that contain a homodimer (chains A and B) bound to a smaller protein (chain C). In this instance, we will call the homodimer the target and the smaller protein will be the binder. To parse this for MAGPIE, the following options can be used:

   ```
   MAGPIE_input_prep.py -i /input -o /output -c A,B -C C
   ```

The -c and -C parameters were chosen because the chains are known and are the same for all the PDB input files.

2. The user has multiple PDB files that contain a homodimer (all different chains) bound to a smaller protein (all different chains). In this instance, we will call the homodimer the target and the smaller protein the binder:

```
MAGPIE_input_prep.py -i /input -o /output -s ARTPGD -S RRRRRRR
```

The -s and -S parameters were chosen as the chains are unknown, but the sequence for the homodimer is the same among inputs ("ARTPGD"), and all the binders contain the sequence "RRRRRRR".

3. The user has multiple PDB files that contain a heterotetramer (all different chains) bound to a smaller protein (chain A). In this instance, we will call the heterotetramer the target and the smaller protein the binder:

```
MAGPIE_input_prep.py -i /input -o /output -s
ARTPGD,GGHTRR,HTWAARR,GGGRRR -C A
```

The -s and -C parameters were chosen as the chains are unknown for the heterotetramer, so four sequences were used. -C was used because the protein binder is always on chain A.

4. The user has multiple PDB files that contain 3 small molecule ligands (LI1, LI2, LI3) bound to a protein target ligand (chain F). In this instance, we will call the small molecules the target and the protein the binder. The user wants to only look at the interactions between LI1 and the target ligand:

```
MAGPIE_input_prep.py -i /input -o /output -L LI1 -C F
```

The -L and -C parameters were chosen as the name is known for the ligand and the chain is known for the binder protein.

5. The user designed some binders *in silico* for an antigen protein. Their goal is to extract all of the binders that bind to the antigen. The chains are all different but the sequences are similar. The binders all start with a FLAG tag while the antigen proteins contain the sequence "DTICIGYHANNSTDTVDTVLEKNV". Here we can treat the antigen as the binder and the binders as the target. The user also wants to align them using the MSA option to get a sequence logo as well.

```
MAGPIE_input_prep.py -i /input -o /output --binder_seqs
DTICIGYHANNSTDTVDTVLEKNV --target_protein_seqs DYKDDDDK --
seq_identity 80 --seq_target_align MSA
```

The --binder_seqs and --target_protein_seqs parameters were chosen because the sequences are known for the target and binders. The --seq_identity option was used because the antigen sequences differ slightly from one another, since they come from different strains. The --seq_target_align option was used to align them via MSA mode.

6. The user has some tetrameric glycoproteins (PFK) with inconsistent chain naming. The user wants to look at the interaction with the ligand named PEP at position 403 within the complex. The user also wants to rename the ligand to chain X and the binders to chain Y.

   ```
   MAGPIE_input_prep.py -i /input -o /output --mesh_search
   ";;PEP;403" --take_first_sm_only --target_sm_3_names PEP --
   search_radius 2 --target_sm_chain_rename X --binder_chain_rename
   Y
   ```

   The --mesh_search was used as we are looking for the subunit in the complex that interacts with only the ligand containing the index 403 and the name PEP. Using the sequence would have found all 4 subunits, thus it cannot be used. The --target_sm_chain_rename and --binder_chain_rename were used as the user preferred to rename them to these identifiers.

**Error and warnings**

1. "Making everything else not the SM the other chain":
   This warning is to ensure the user understands that all other proteins are binders. This occurs when the user does not specify a binder chain or sequence and only a small molecule.

2. "ERROR no chain or seq selected for binder"
   This error occurs when no binder, small molecule, or target protein is specified.

3. "ERROR no chain or seq selected for target_protein"
   This error occurs when no target ligand information is specified. This error is unlikely to occur, because another check should detect this first.

4. "You must enter a target protein chain and/or sequence or small molecule name and/or chain or both!"
   This error occurs when no target ligand information is specified.

5. "the input path you entered does not exist!"
   This error occurs when the pdb file or directory with the -i parameter is wrong.

6. "the output path you entered does not exist!"
   This error occurs when the output directory with the -o parameter is wrong.

7. "the binder or target seq fa file does not exist"
   This error occurs when the .fasta file for the binder or target does not exist.

8. "the binder or target seq fa file is a directory and not a file"
   This error occurs when the .fasta file for the binder or target is a directory and not a file.

9. "the reference sm file does not exist!"
   This error occurs when the PDB file for the small molecule reference does not exist.

10. "the reference sm file given was a directory not a file!"
    This error occurs when the PDB input for the small molecule target ligand reference structure is a directory and not a file.

11. "the reference sequence alignment file does not exist!"
    This error occurs when the PDB reference for sequence alignment does not exist.

12. "the reference sequence alignment file given was a directory not a file!"
    This error occurs when the PDB reference for sequence alignment is a directory and not a file.

13. "You must use target_sm_index with either target_sm_chains or target_sm_3_names"
    This error occurs when the user uses the index option when no chain or name for the small molecule is set.

14. "seq_target_ref_pdb cannot be used with only one pdb file"
    This error occurs when the user specifies a target reference PDB for sequence alignment and is using a single PDB file as input.

15. "non .pdb file types are not supported!"
    This error occurs when the user has file types other than .pdb in the input directory.

### 7. Multithreading and performance benchmarks

When using larger datasets, the Google CoLab version is not optimized to parallellize threads and can increase MAGPIE compute time. Within the local version, MAGPIE allows users to multi-thread using CPUs to efficiently run larger datasets. Users only need to run cell **3.1 Advanced Options Multithreaded** and specify the number of threads to use.

We tested the runtime of the MAGPIE local installation on multiple machines during the rate-limiting 3D plotting step on the COA small molecule ligand example, Conformer Pool 11 ( containing 27 models), the SARS-CoV-2 spike protein example,and 1570 PDBs generated by expanding the Protein Example dataset. The distance option was set to 8 Å, default options for clustering were used, and time of execution was recorded in seconds.

<u>Processor: 12th Gen Intel(R) Core(TM) i7-12700H 2.30 GHz</u>
<u>Installed RAM 24.0 GB (23.7 GB usable)</u>

| **Example→ N threads ↓** | **Small molecule ligand example, reference_11** | **Protein example** | **1570 PDBs** |
|---|---|---|---|
| 1 | 10.7 s | 10.2 s | 217.6 s |
| 2 | 9.0 s | 5.9 s | 118.9 s |
| 4 | 6.5 s | 4.7 s | 73.1 s |
| 8 | 2.6 s | 2.9 s | 53.6 s |
| 16 | 2.3 s | 2.5 s | 45.2 s |

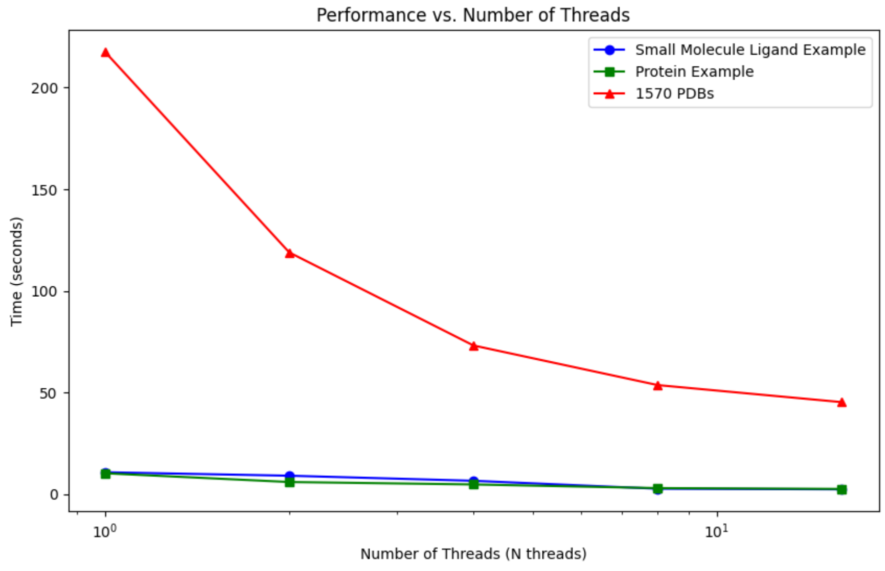

Processor: AMD Ryzen 9 7950X
Installed RAM 64.0 GB (63.1GB usable)

| **Example→ N threads ↓** | **Small molecule ligand example, reference_11** | **Protein example** | **1570 PDBs** |
|---|---|---|---|
| 1 | 7.8 s | 7.2 s | 172.2 s |
| 2 | 4.7 s | 4.0 s | 98.9 s |
| 4 | 2.8 s | 2.4 s | 77.82 s |
| 8 | 2.1 s | 1.4 s | 26.4 s |
| 16 | 1.0 s | 0.95 s | 16.2 s |

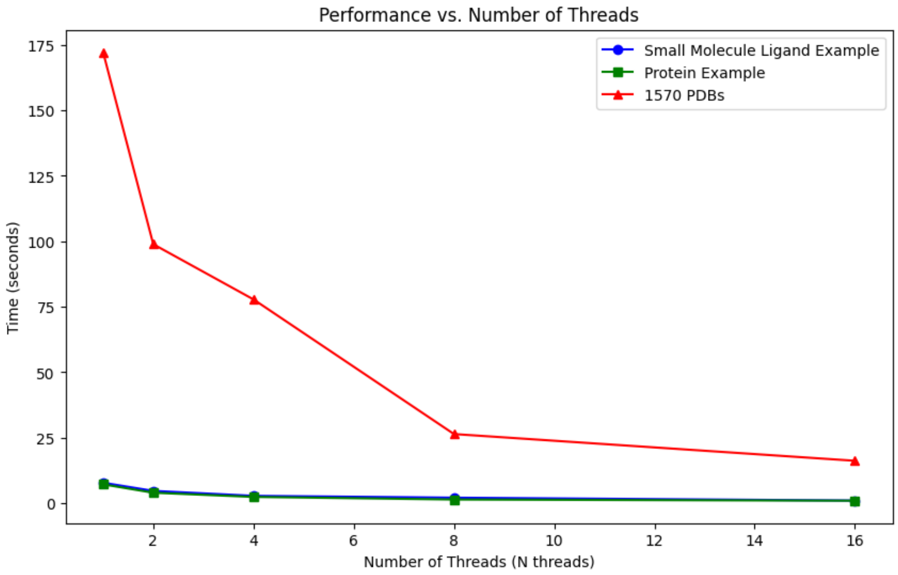

### 8. Additional information about protein complexes in Figures 2-6

**Case study #1: the target ligand is a protein**

We used 63 complexes of antibody Fabs bound to the SARS-CoV-2 spike protein receptor binding domain (RBD) from the CoV3D database.[11] These were globally aligned on the spike RBD in PyMOL using the "align" command, and then all structures were exported as individual PDB files. The PDB codes corresponding to the structures are: 7l5b, 7k8z, 7kn5, 7k4n, 7k9z, 7k90, 7cai, 7jv6, 7dk4, 6yz5, 7jx3, 7k9z, 7c01, 6zh9, 7k43, 7bz5, 6xc3, 7kmi, 7jx3, 7chb, 6xc4, 7kmh, 7jx3, 7chf, 7jmw, 7kmg, 7c8v, 7chf, 7jmo, 7c8w, 7b3o, 7ch5, 7jmp, 7jvb, 7kgk, 7byr, 7kzb, 7cwn, 7a29, 7k8t, 6yla, 7bwj, 7kgj, 7k8m, 7cm4, 7cdj, 7klw, 7k8u, 6xkq, 7cdi, 6zxn, 6xcm, 6xkp, 7cjf, 7kn5, 7k8v, 6xe1, 6xdg, 7kn6, 7k8w, 7ld1, 6xdg, 7kn7, 7k8x, 6zcz, 7jvc, and 7kkl.

**Case study #2: the target ligand is a small molecule**

We used 199 structurally diverse bacterial enzymes that bind coenzyme A (COA). We searched the PDB for structural models with refinement resolutions between 1.5 and 3 Å using its PubChem identifier code 87642. From this set of >600 structures, to reduce redundancy and noise in the dataset, we chose 199 models randomly. Using MAGPIE_input_prep.py with the small molecule target ligand name and mesh area search selection options, we removed all other chains that were not COA or the protein(s) bound/nearby to COA, including redundant protein and COA chains.

<u>Cleaning command line:</u>

```
python ~/MAGPIE/MAGPIE_input_prep.py -i <input_directory> -o <output_directory> -L COA -M 'A,B;;COA;'
```

The structures were then globally aligned on COA using align_small_molecule.py, and then all structures were exported as individual PDB files.

<u>Alignment command line:</u>

```
python ~/MAGPIE/align_small_molecule.py -c B -T 2.5 -i <input_directory> -o <output_directory> -p True
```

The PDB codes corresponding to the structures are: 1m4g, 1vpm, 3ddd, 5szu, 8b32, 1p0h, 2eis, 3fbu, 5tvj, 8b3c, 1pu9, 2pfr, 4ien, 6wfk, 1pua, 2v1o, 5gi6, 6ygd, 1q2d, 3b7k, 5kl9, 7cz3, 1jll, 2scu, 4yak, 7lcl, 8ciw, 1qfl, 2vtz, 6aqp, 7ld2, 2f2s, 2wkt, 6bja, 7ldc, 2hqy, 4xyl, 6pfn, 7ldu, 2nu7, 4xz3, 7cw5, 7mss, 3s6g, 4hzo, 5hwp, 6j1i, 7ztk, 1esm, 3wy0, 4jd3, 2zsd, 4ag9, 6ktq, 1h1t, 1t4c, 2qf7, 3ubm, 6v8k, 1q6y, 1xvt, 2vjl, 6cz6, 2ahv, 4eu6, 4eud, 2onf, 4eub, 8i40, 1yqz, 2vfc, 4eqs, 5szy, 6gzt, 2c43, 4fc7, 5xxs, 7bcz, 7yjb, 2wdo, 4qjl, 6lq4, 7ed1, 3hqj, 4r3k, 6rcx, 7n8m, 3wd7, 5ahs, 6wf3, 7yj6, 2rkv, 2zw7, 3cgc, 4fx9, 6ruz, 2zw4, 3b2s, 3ict, 5ktd, 3p3i, 8hjb, 1ixe, 3fsb,

5vxc, 7ciw, 8dqr, 1n8w, 3pvy, 6as5, 7n0l, 8pn7, 2cts, 3rq5, 6boo, 7o4s, 8u2t, 2gq3, 3vbk, 6j1e, 7pyt, 2req, 3vbm, 6lpv, 7rkz, 3cts, 4l9z, 6wqc, 7rmp, 2d5a, 2v18, 2yj0, 1bo4, 2p8u, 6vr2, 7zkt, 2b58, 5us1, 7kps, 1pg3, 4r1l, 8p5u, 1ebl, 5iv0, 2h3p, 3otw, 5yh7, 2h3u, 4x0o, 5yrr, 3pzc, 5kp2, 4psx, 1n71, 4u9w, 6bvc, 6wfg, 7wx6, 2cnt, 5f48, 6ksb, 6ygb, 2prb, 5hmn, 6lq1, 7csj, 3owc, 5n1u, 6lq8, 7q3a, 1dqa, 4m20, 6he2, 3lbe, 4rvn, 8i8m, 1eab, 7uuo, 7yxm, 2gyo, 4hzp, 4nhd, 2d3m, 4b3j, 5o9v, 6ehj, 4b3i, 5o9u, 5vj1, 6p5u, 5ht0, 6mb9, 7mqk, 1h16, 8gju, 4kec, 6wuk, 7fc7.

**Case study #3**

468 PFK-1 representative UniRef90 sequences were selected after MMseqs2-based clustering[12] of a multi-sequence alignment with *Escherichia coli* PFK-1 as the query sequence. We took UniRef90 sequences corresponding to the top 7 phyla (n=251) and mapped them to UniProtKB ID's using the online UniProt mapping tool. Mapping from UniRef90 to UniProtKB ID was done to retrieve the full amino acid sequence of each entry. This is because the sequence found in the MMseqs2 MSA output only represents the portion of the sequence that is aligned in the MSA. Next, these UniProtKB sequences were filtered to exclude sequences with duplicate taxonomical identifier numbers (OX number), lack of taxonomical designation, and high sequence identity (>60%), resulting in 83 UniProtKB sequences from 7 phyla and 42 orders which were used as input for structural prediction using AlphaFold2.[13] PEP was subsequently modeled into the allosteric pocket of each structural model by structural alignment with the previously solved mutant *Geobacillus stearothermophilus* PFK-1-PEP (PDB: 4I4I) complex.[14] Next, the TM-score and RMSD were calculated via USalign[15] and outliers with an RMSD greater than 4.5 Å (UniProt ID's A0A9D1NFY2 and A0A1I5YU75) and a TM-score[16] below 0.7 (UniProt IDs: A0A1I5YU75, also an RMSD outlier) were excluded. The ligand total energy was then calculated via Rosetta scoring[17] and models with positive energies were excluded (A0A2W4FR96 and A0A9E3QUE3)**.** For reference, the average ligand total energy for the PEP analog 2-phosphoglycolic acid (PGA) in our relaxed 6PFK structure is -1.69 REU, and -2.215 for our relaxed 4I4I structure. Models with the ligand bound outside of the binding pocket, which was evaluated via PyMol visualization of each structure, were also excluded. After these final filtering steps, 72 PEP-bound PFK-1 models from 7 phyla and 39 orders were used as input for MAGPIE visualization (Figure 5).

Using MAGPIE_input_prep.py with the small molecule target ligand name and residue index selection options, we removed all but one PEP molecule, and subsequently used MAGPIE_align_small_molecule.py with a 0.4 Å global RMSD threshold to bin PEP conformers.

Based on the 83 AlphaFolded structural models used in the PEP-bound PFK-1 MAGPIE analysis, ADP was modeled into the allosteric pocket of each structural model in the same way, using the previously solved *Escherichia coli* PFK-1-ADP (PDB: 1PFK) complex as a reference (Figure 5). After applying the filtering protocol described previously, 65 PFK-1 sequences from 4 phyla and 15 orders were used as input for MAGPIE visualization. Outliers with an RMSD greater than 4.5 Å (UniProt ID's A0A9D1NFY2 and A0A1I5YU75) and a TM-score below 0.7 (UniProt IDs: A0A1I5YU75, also an RMSD outlier, and A0A350LL64) were excluded. Structures with positive total ADP score were excluded (I9NPE1, A0A9C7AUT3, A0A2W4FR96, A0A255E815, A0A2M8PW24, A0A6N7V2J5, A0A926S2G2, A0A8G1T9F0, A0A1P8UHZ0, M2U2S2, L0IMU1). The average sequence identity within each phyla was 54.9% and the average sequence identity within any two phyla was 25.98%.

Cleaning command line:
PFK-1-PEP
python ~/MAGPIE/MAGPIE_input_prep.py -i <input_directory> -o <output_directory> -L PEP -I 401
PFK-1-ADP
python ~/MAGPIE/MAGPIE_input_prep.py -i <input_directory> -o <output_directory> -L ADP -I 401

Alignment command line:
PFK-1-PEP
python ~/MAGPIE/align_small_molecule.py -c B -T 0.4 -i <input_directory> -o <output_directory>

PFK-1-ADP
python ~/MAGPIE/align_small_molecule.py -c B -T 0.75 -i <input_directory> -o <output_directory>

The UniProt IDs corresponding to the PEP-bound structures for are: A0A1B7KSA1, A0A7X2PK64, A0A6N7EXS0, A0A9D9EWD2, A0A959ZK87, A0A2D9T5C8, A0A9D5T0S7, A0A1G9V4P9, A0A347WIF3, A0A926S2G2, M2U2S2, A0A143DEI8, A0A9C7AUT3, A0A926K9E6, A0A2U1TK21, A0A949N606, A0A4R6N9I0, A0A9E3QUE3, A0A0M8K9I8, A0A522A2V8, A0A1I5PL87, A0A356DVB6, A0A6P2BVL8, A0A845QDK5, A0A117S2W8, A0A931J453, A0A954S093, A0A849K9K2, I8HY23, A0A1H8ZQQ9, A0A317MRI0, A0A7L9RTV1, A0A960DEI3, B1Y8G4, A0A255E815, A0A2M8PW24, A0A953BG47, A0A7W2T3Z8, L0IMU1, A0A1T4LYW8, A0A1Y4URI3, A0A9D1DYX9, A0A0P6WUL9, A0A8G1T9F0, A0A2Y9CKF7, A0A7W1TMQ5, A0A0K1P8E0, A0A6N7V2J5, A0A0P6YHQ1, A0A0P9EST4, A0A2W4FR96, A0A5C6B571, A0A0F6YL10, A0A4Y8ZXG8, A0A6J4Q7L3, A0A9D1CHV2, A0A957EMT2, A0A1P8UHZ0, A0A940J4H3, M5EFB6, A0A7Y6UDE6, I9NPE1, A0A2N3Q0X7, A0A9D6WYZ8, A0A085W4P3, A0A357ARF4, A0A419GUH8, A0A3L7Y257, R7K4H7, A0A7Y6YDF6, A0A6B1B3N7, A0A7Y5RS15, A0A1G9NID4, A0A350LL64

The UniProt IDs corresponding to the ADP-bound structures for are: A0A3L7Y257, A0A4V2AJ23, A0A0F6YL10, A0A4R6N9I0, A0A957EMT2, A0A1C6JNA4, A0A6J4Q7L3, A0A2D9T5C8, A0A2N3Q0X7, A0A6P2BVL8, A0A143DEI8, A0A7Y5RS15, A0A954S093, A0A085W4P3, M5EFB6, A0A9D6WYZ8, A0A0P6WUL9, A0A1H8ZQQ9, A0A1Y4URI3, R7K4H7, A0A1G9V4P9, A0A7Y6YDF6, A0A077AYE3, A0A926K9E6, A0A2Y9CKF7, A0A1B7KSA1, A0A7X2PK64, A0A356DVB6, A0A419GUH8, A0A357ARF4, A0A0P9EST4, A0A0M8K9I8, A0A1T4LYW8, A0A9D5T0S7, A0A357KWS7, A0A949N606, A0A931J453, A0A7L9RTV1, A0A953BG47, A0A6N7EXS0, A0A0P6YHQ1, A0A849K9K2, A0A0K1P8E0, A0A845QDK5, A0A522A2V8, A0A5C6B571, A0A960DEI3, A0A1G9NID4, A0A9E3QUE3, A0A6B1B3N7, A0A317MRI0, A0A9D1CHV2, A0A940J4H3, A0A117S2W8, A0A9D9EWD2, A0A7W1TMQ5, B1Y8G4, A0A7W2T3Z8, A0A1I5PL87, A0A2U1TK21, A0A7Y6UDE6, A0A4Y8ZXG8, A0A9D1DYX9, A0A347WIF3, A0A959ZK87

### 9. Local installation requirements

MAGPIE requires the following Python packages and versions to run the local version:
Jupyter (1.0.0)
Numpy (1.18.5)
Pandas (1.3.4)
Matplotlib (3.4.3)
Glob (0.7)
Plotly (5.9.0)
Spicy (1.7.1)
Pymol (2.5.8)
Seaborn (0.12.2)
Muscle (5.1)

### 10. Example Conda environment install

create create MAGPIE
conda activate MAGPIE
conda install -c conda-forge -c schrodinger pymol-bundle
conda install conda-forge::biopython
conda install bioconda::muscle
conda install conda-forge::plotly
conda install conda-forge::glob2
python -m pip install scipy
conda install conda-forge::matplotlib
conda install bioconda::logomaker
conda install anaconda::seaborn
pip install -U scikit-learn

### 11. References

1. Cock PJA, Antao T, Chang JT, Chapman BA, Cox CJ, Dalke A, Friedberg I, Hamelryck T, Kauff F, Wilczynski B, et al. (2009) Biopython: freely available Python tools for computational molecular biology and bioinformatics. Bioinformatics 25:1422–1423.

2. Hasegawa M, Noda H (1975) Distribution of hydrogen bond angles in molecular crystals. Nature 254:212–212.

3. Kurczab R, Śliwa P, Rataj K, Kafel R, Bojarski AJ (2018) Salt Bridge in Ligand–Protein Complexes—Systematic Theoretical and Statistical Investigations. J. Chem. Inf. Model. 58:2224–2238.

4. Berman HM, Westbrook J, Feng Z, Gilliland G, Bhat TN, Weissig H, Shindyalov IN, Bourne PE (2000) The Protein Data Bank. Nucleic Acids Research 28:235–242.

5. Evans R, O'Neill M, Pritzel A, Antropova N, Senior A, Green T, Žídek A, Bates R, Blackwell S, Yim J, et al. (2022) Protein complex prediction with AlphaFold-Multimer. :2021.10.04.463034. Available from: https://www.biorxiv.org/content/10.1101/2021.10.04.463034v2

6. Gray JJ, Moughon S, Wang C, Schueler-Furman O, Kuhlman B, Rohl CA, Baker D (2003) Protein–Protein Docking with Simultaneous Optimization of Rigid-body Displacement and Side-chain Conformations. Journal of Molecular Biology 331:281–299.

7. Anon The PyMOL Molecular Graphics System, Version 2.0 Schrödinger, LLC.

8. Pettersen EF, Goddard TD, Huang CC, Meng EC, Couch GS, Croll TI, Morris JH, Ferrin TE (2021) UCSF ChimeraX: Structure visualization for researchers, educators, and developers. Protein Science 30:70–82.

9. Pedregosa F, Varoquaux G, Gramfort A, Michel V, Thirion B, Grisel O, Blondel M, Prettenhofer P, Weiss R, Dubourg V, et al. (2011) Scikit-learn: Machine Learning in Python. Journal of Machine Learning Research 12:2825–2830.

10. Ester M, Kriegel H-P, Sander J, Xu X A density-based algorithm for discovering clusters in large spatial databases with noise. In: Proceedings of the Second International Conference on Knowledge Discovery and Data Mining. KDD'96. Portland, Oregon: AAAI Press; 1996. pp. 226–231.

11. Gowthaman R, Guest JD, Yin R, Adolf-Bryfogle J, Schief WR, Pierce BG (2021) CoV3D: a database of high resolution coronavirus protein structures. Nucleic Acids Research 49:D282–D287.

12. Steinegger M, Söding J (2017) MMseqs2 enables sensitive protein sequence searching for the analysis of massive data sets. Nat Biotechnol 35:1026–1028.

13. Jumper J, Evans R, Pritzel A, Green T, Figurnov M, Ronneberger O, Tunyasuvunakool K, Bates R, Žídek A, Potapenko A, et al. (2021) Highly accurate protein structure prediction with AlphaFold. Nature 596:583–589.

14. Mosser R, Reddy MCM, Bruning JB, Sacchettini JC, Reinhart GD (2013) Redefining the Role of the Quaternary Shift in *Bacillus stearothermophilus* Phosphofructokinase. Biochemistry 52:5421–5429.

15. Zhang C, Shine M, Pyle AM, Zhang Y (2022) US-align: universal structure alignments of proteins, nucleic acids, and macromolecular complexes. Nat Methods 19:1109–1115.

16. Zhang Y, Skolnick J (2005) TM-align: a protein structure alignment algorithm based on the TM-score. Nucleic Acids Research 33:2302–2309.

17. Alford RF, Leaver-Fay A, Jeliazkov JR, O'Meara MJ, DiMaio FP, Park H, Shapovalov MV, Renfrew PD, Mulligan VK, Kappel K, et al. (2017) The Rosetta All-Atom Energy Function for Macromolecular Modeling and Design. J. Chem. Theory Comput. 13:3031–3048.
